# Colonial legacies in eponymous species names: a global network perspective

**DOI:** 10.64898/2026.07.25.740734

**Authors:** Akshay Bharadwaj, Joel L. Pick, Bernd Lenzner

## Abstract

Eponymous etymologies (i.e., organisms named after people) are prevalent in global taxonomy worldwide, across scientific and common names. The persistent usage of eponyms has been intensely debated in recent times, across various sociopolitical, applied and philosophical perspectives. However, a rigorous statistical analysis of eponyms and their enduring colonial legacies remains hitherto unexplored. Here, we use a global and spatially-explicit network analysis to examine 2416 contemporary bird eponyms (21% of all bird species on earth) and their relationships with historical European colonialism. Additionally, we develop and utilise a Colonial Exposure Index (CEI) that summarizes the geopolitical influence of various colonial empires on modern-day countries. Our results show that an overwhelming majority of bird eponyms worldwide are named after people belonging to countries where the bird species does not occur. A majority of these eponyms honor scientists, aristocrats and army officers from erstwhile colonial empires, with eponymous species descriptions peaking during colonial expansion. Importantly, our analysis highlights a positive relationship between the Colonial Exposure Index (a metric factoring the duration and area of a country colonized by a specific empire) and the number of eponymous bird species honoring persons from that specific empire. Our temporal analysis, however, shows that the use of eponyms has recently shifted towards honoring native persons and distinguished conservation champions. Overall, this study presents an important statistical examination of eponyms and their colonial legacies, thereby providing valuable context for ongoing debates on the reform or retention of eponyms worldwide.

## Introduction

Science is inextricably linked to the prevailing sociopolitical context [1,2]. Following the late 15th century ‘Age of Exploration’, European colonialism emerged as the dominant global force expanding trade and transport through the exploitation of distant lands and/or peoples [3,4]. This expansion facilitated the spread of Western scientific knowledge systems, including the Linnean taxonomic classification, which remains foundational today [5]. Following nomenclatural conventions, the name of a species new to Western science is chosen and published by the first describer [6]. Several species names (particularly in western Europe) stem from ancient Anglosaxon origins often predating Linnean taxonomy, such as the Common Cuckoo *Cuculus canorus* [7] and Common Nightingale *Luscinia megarhynchos* [8]. Other species names contain information on notable morphological features (e.g., White-collared Blackbird *Turdus albocinctus*) or the geographic range of the species (e.g., Himalayan Griffon *Gyps himalayensis*). However, a longstanding and common practice is the first describer naming a new species after people – such as their family members (e.g., Mrs. Gould’s Sunbird *Aethopyga gouldiae*), patrons of expeditions (e.g., Frances’s Sparrowhawk *Tachyspiza francesiae*), their scientific colleagues (e.g., Archbold’s Nightjar *Eurostopodus archboldi*) or other influential figures (Victoria Crown Pigeon *Goura victoria*). Such species names which contain the names of human beings are referred to as ‘eponyms’.

The use and persistence of eponyms has generated substantial debate recently across various scientific fields, ranging from medical science [9,10], applied physics and engineering, [11] and the taxonomy of birds [12], primates [13], molluscs [14], amphibians [15] and reptiles [16]. This debate on the retention or reform of eponyms has widespread consequences for global scientific research, particularly in biodiversity science and taxonomy [17–25]. Presently, nomenclatural changes are reviewed and/or revised by specialized committees on a case-by-case basis. For example, the American Ornithological Society (AOS) recently declared its push to remove eponyms from the common names of American birds altogether [26]. At the same time, we observe more eponyms honoring local persons [e.g., the Munchique Wood-wren Henicorhina negreti from the Colombian Andes honoring the Colombian scientist Álvaro José Negret, 27]. Nevertheless, eponyms honoring non-native persons remain widespread in global taxonomy today.

Critics argue that eponyms are an enduring honorific reminder of former benefactors of institutions with racist or colonialist roots, often ignoring indigenous knowledge and local naming traditions [19]. Concerns related to Diversity, Equity and Inclusion in modern science [DEI; 28] highlight the current lack of acknowledgement and credit for local, indigenous and historically underrepresented groups. Even within post-colonial taxonomy (i.e., since 1950), studies show only a minimal albeit increasing representation of Global South persons honored in eponymous descriptions, despite most species descriptions occurring in the Global South [12,14]. Consequently, several scientists have called for banning eponyms altogether, calling their persistent usage “unnecessary and objectively difficult to justify” [19]. The abolishment of eponyms is also seen part of broader efforts to discourage parachute science, i.e., scientists and funders from the Global North setting research agendas and getting credit for studies conducted in the Global South, which often still follow colonial legacies [14,29–31].

However, not all scientists are in agreement. Others argue that the abandonment of eponyms threatens nomenclatural stability [17,but see 25,32], impedes taxonomical research, and only represents virtue signaling by researchers from the Global North [24]. Pethiyagoda (2023) argues that such changes would compel taxonomists in biodiverse regions, especially in developing countries, to direct their attention away from combating the rapidly growing threats to biodiversity itself [33]. Furthermore, Jiménez-Mejías et al. (2024) caution that sweeping taxonomic changes removing eponyms would threaten the value of nomenclature codes, specifically its universality and stability [34]. Moreover, recent eponyms increasingly acknowledge local scientists, indigenous leaders, women and non-western honorifics [27,35]. Some also note that this practice helps to raise awareness and funds for conservation work, logistics, as well as research and training of taxonomists in regions with fewer financial resources [18,20].

Between these two opposing opinions, several middle-ground solutions have been proposed. For example, Thiele (2023) suggests that the eponyms honoring people who participated in clearly egregious actions such as slavery, mass murder or acts of significant colonial injustice are highly problematic and need urgent renaming. [24]. Critical cases could, for example, be reviewed by a specialized committee with well-defined procedures to remove overtly offensive or derogatory names [23]. A recent example of such a process was the proposal by Smith & Figueiredo (2021) to “permanently and retroactively eliminate epithets with the root *caf[e]r-* or *caff[e]r*- from the nomenclature of algae, fungi and plants” that was adopted at the International Botanical Congress in Madrid [36].

Recent heated debates on eponyms, whether advocating reform or retention, have primarily focused on case studies and opinion pieces. Few studies examine the phenomenon quantitatively. DuBay et al. (2020) report that 95% of the scientific names of bird species described since 1950 occurred in the Global South, although they disproportionately honored individuals from the Global North. Notably, in recent times, there is an increase in eponyms honoring persons from the Global South [27] and in descriptions co-authored by Global South persons [12]. Notwithstanding its global relevance, a formal statistical examination of the phenomenon, i.e., the relationship between eponym usage today and its colonial legacies, remains hitherto unexplored.

Here, we present a spatially-explicit, statistical assessment of the extent of colonial legacies in bird eponyms today, across both scientific and English common names. We investigate eponym occurrence and frequency in relation to the geographic area and duration of colonial occupation by European colonial powers. We hypothesized *a priori* that (a) most eponyms honor non-native persons (largely from former empires), and (b) the number of eponymous birds honoring individuals from a given empire is strongly associated with that empire’s historical influence on the country.

To do so, we combine bird eponym datasets with the incidence and extent of colonial occupation by the eight major European empires. Specifically, we introduce a Colonial Exposure Index (CEI) that incorporates information on the duration of colonial occupation and its spatial extent within each modern-day country. Using bipartite networks [37,38], for each country, we examine relationships between the level of exposure to each colonial empire and the number of bird eponyms honoring persons from that specific empire [39]. We further investigate whether this relationship differs between scientific and common names, assuming that common names are more flexible when compared to scientific names that are regulated by the International Code of Zoological Nomenclature (ICZN) and subject to rigid rules that prioritize the oldest correctly published name [40,41]. Although eponyms exist across all taxa, we focus on birds because they are one of the most accessible taxonomic groups, and often a gateway to exploring the natural world for many [42]. Our statistical examination of colonial legacies in eponyms today contextualizes and facilitates informed discussions among researchers and civil society at large.

## Results

### Spatio-temporal (mis-)match between origin of bird eponym and its namesake

Of the 11,195 total bird species in the world, 2,416 species (21%) are eponyms. The highest numbers of bird eponyms were found in Indonesia (n = 288), followed by Peru (n = 285) and Colombia (n = 270). Madagascar (20.6%) showed the highest relative numbers of bird eponyms in relation to its total species richness, followed by the Comoros (19.8%) and Republic of Congo (19.2%; Table S1). An overwhelming majority of bird eponyms honor persons who are not from the countries where the bird species is found (observe dominance of grey lines in Fig 1a). Most eponyms were attributed to people from Great Britain (30% of scientific and 33% of common names; Table S2), followed by France (16% and 8% respectively; Table S2) and Germany (15% and 16% respectively; Table S2). Overall, results across scientific names and common names showed very similar patterns (Fig S3-S5). With regard to country-matching eponyms (i.e., where the native country of the bird coincides with that of its namesake; Table S3) the USA stood out (n = 41), distantly followed by South Africa (n = 6) and New Zealand (n = 4).

**Figure 1:**
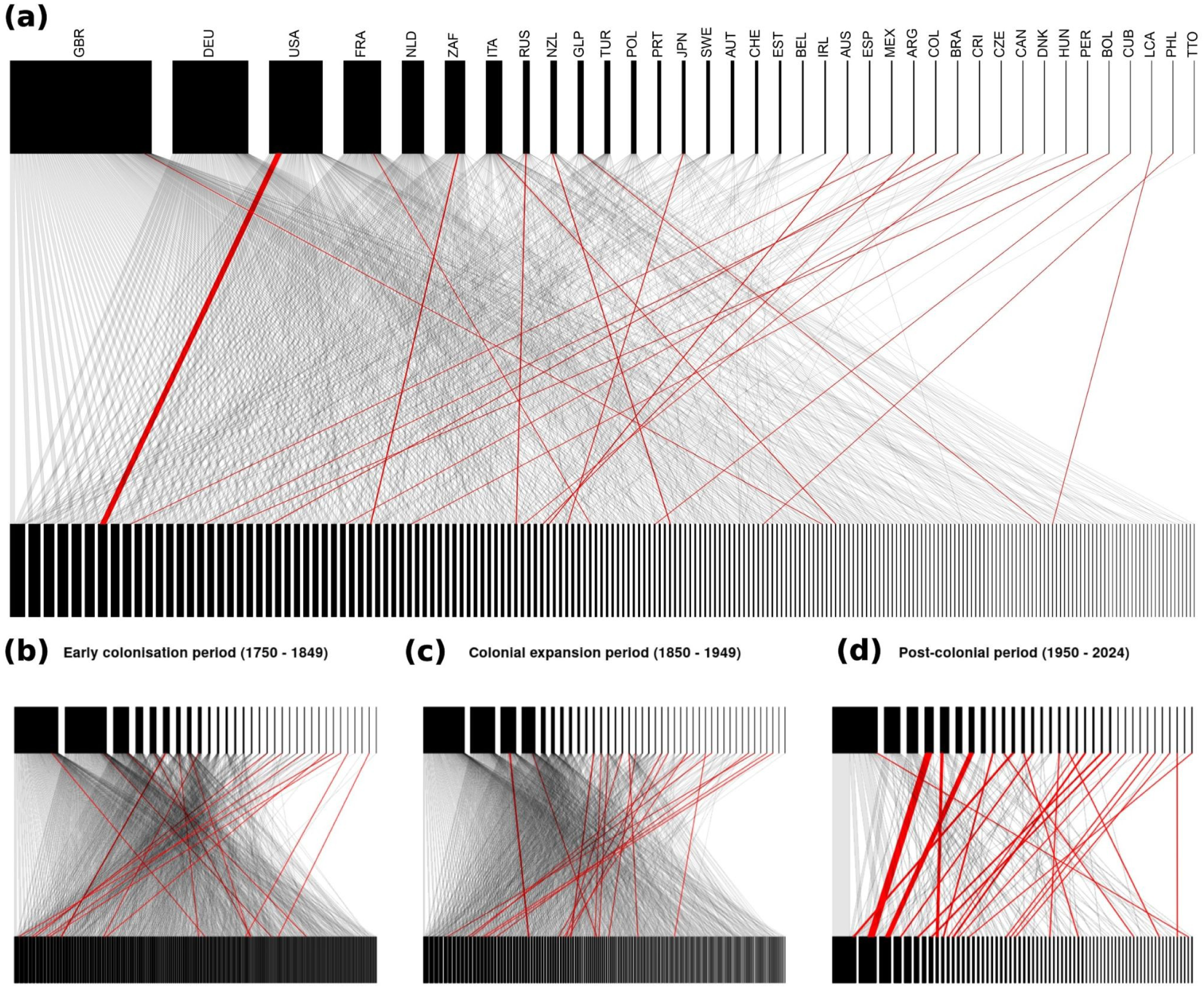
Bipartite networks visualizing relationships in eponymous bird scientific names (a) overall, (b) in the early colonization period, (c) colonial expansion period and (d) post-colonial period. In each network, countries where eponymous birds occur are shown in the bottom layer, while the country of origin of the persons honored in the eponymous bird name is shown on the top layer. A gray edge between the layers denotes a single eponymous bird species found in a country when it is named after a person who is not from that country. Conversely, red edges highlight cases where the bird and the person honored in its name are from the same country. Early colonial and colonial expansion periods are dominated predominantly by non-native eponym descriptions (networks dominated by gray edges). The proportion of red edges relative to the gray, increases in the post-colonial period, in line with findings from Sangster (2025). The full key to all ISO country codes is provided in Table S1.

To understand the temporal dynamics of the bipartite network structure, we subdivided the data into 3 distinct time periods: (i) the early colonization period (1750 – 1849; Fig 1b), (ii) the colonial expansion period (1850 – 1949, Fig 1c), and the post-colonial period (1950 – 2024; Fig. 1d). In the early colonization period, only 9.8% (n = 45) were country-matching eponyms, falling to 8.8% (n = 134) in the colonial expansion period, and climbing to 32.6% (n = 61) in the post-colonial period. This trend shows that while overall new bird species descriptions are declining, the share of country-matching eponyms is increasing (as is observable by the increasing proportion of red edges in Fig 1d). These time windows were chosen *a priori* and are independent of the colonial legacy of any one country; rather, they are specifically to examine how above-mentioned relationships change over time.

Investigating the temporal patterns in eponym description further, we found a clear increase in the number of eponyms particularly honoring persons from European colonial powers with the onset of the 19^th^ century. This increase tapers off close in the first half of the 20^th^ century (Fig 2A). In terms of their professions, most eponyms today honor scientists (40.4%), followed by naturalists and army personnel (17.8% and 11.5% respectively). Other occupations are only represented by a few eponyms (see Fig 2B and Table S4).

**Figure 2:**
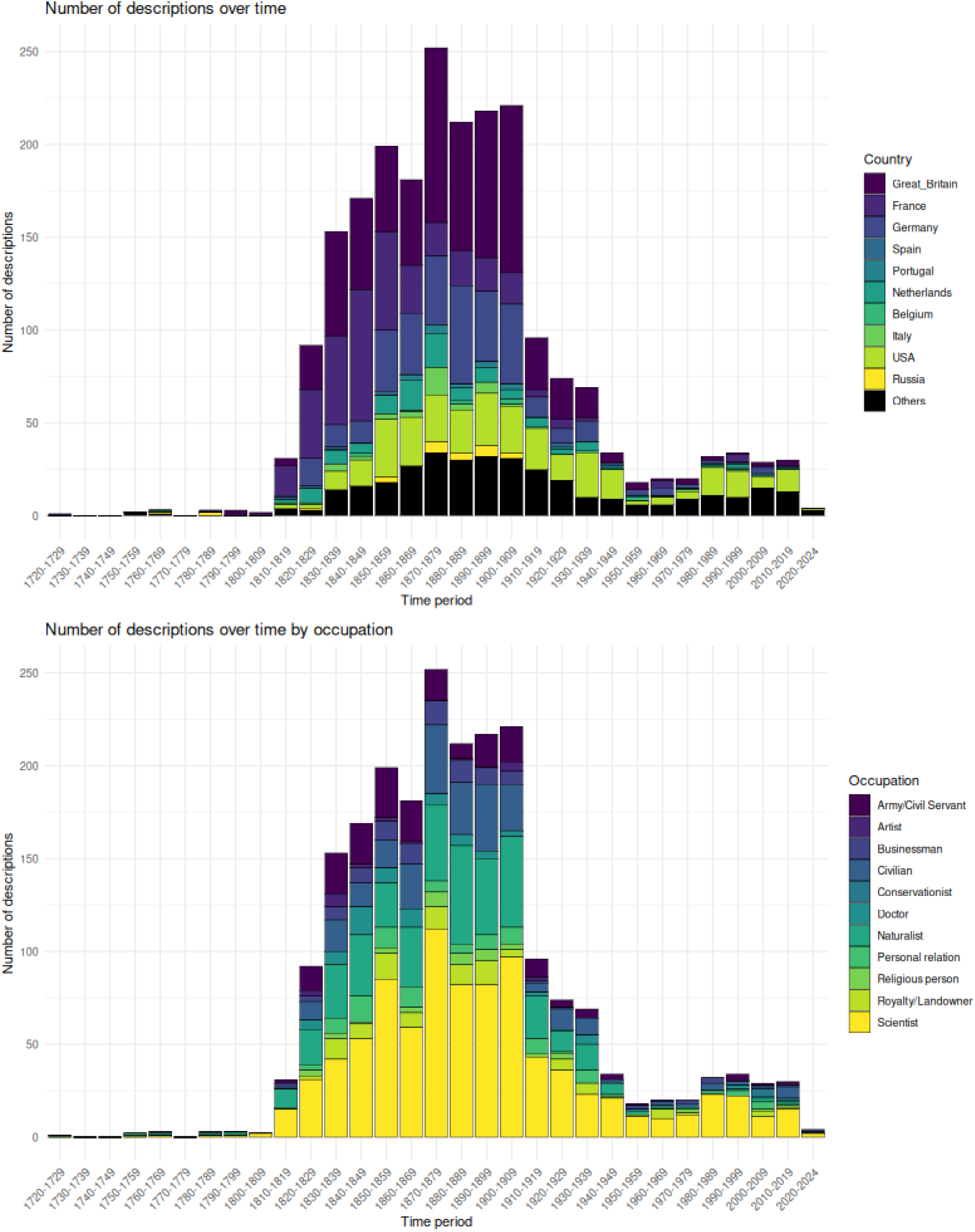
**(A)** Number of new eponymous bird descriptions per decade, shown as stacked bars with colors indicating the nationality of the individuals honored. **(B)** Number of new eponymous bird descriptions per decade, shown as stacked bars with colors indicating the professions of/reasons behind the individuals being honored.

### European colonial legacies in bird eponyms

We further analyzed the lasting legacies of European colonialism in eponyms today. Specifically, we analyzed patterns in the 1993 eponyms (a major subset of the global data) which honor people from the eight European empires, as they relate to patterns in the Colonial Exposure Indices.

Overall, in both the colonialism and eponym bipartite networks, Great Britain dominates the network in the sheer number of eponyms honoring its citizens, followed by France and Germany (Fig 3). Given the large number of colonizer-country combinations that had zero eponyms linking them, we tested the effect of colonialism on eponym counts using a Zero-inflated Poisson model, which explicitly separates (a) the Zero-inflated component: effect of CEI on the probability of having *any* eponyms; and (b) the Poisson component: effect of CEI on the number of eponyms in the cases where there is some probability of having them (expected eponym count greater than zero). In line with our hypothesis we found a positive relationship between CEI and the number of eponyms, when there was some probability of having them, i.e., in the Poisson component. This relationship was consistent across scientific names (Table 1) and common names (Table S6). Furthermore, the probability of having any eponyms increased with increasing CEI, as indicated in the Zero-inflated component of the model (although the 95% credible intervals did include zero here). We further tested the statistical support for an overall effect of CEI (i.e. in both components combined), and found a positive effect of CEI on the number of eponyms in >99% of iterations. All model outputs are shown in Table 1.

**Figure 3:**
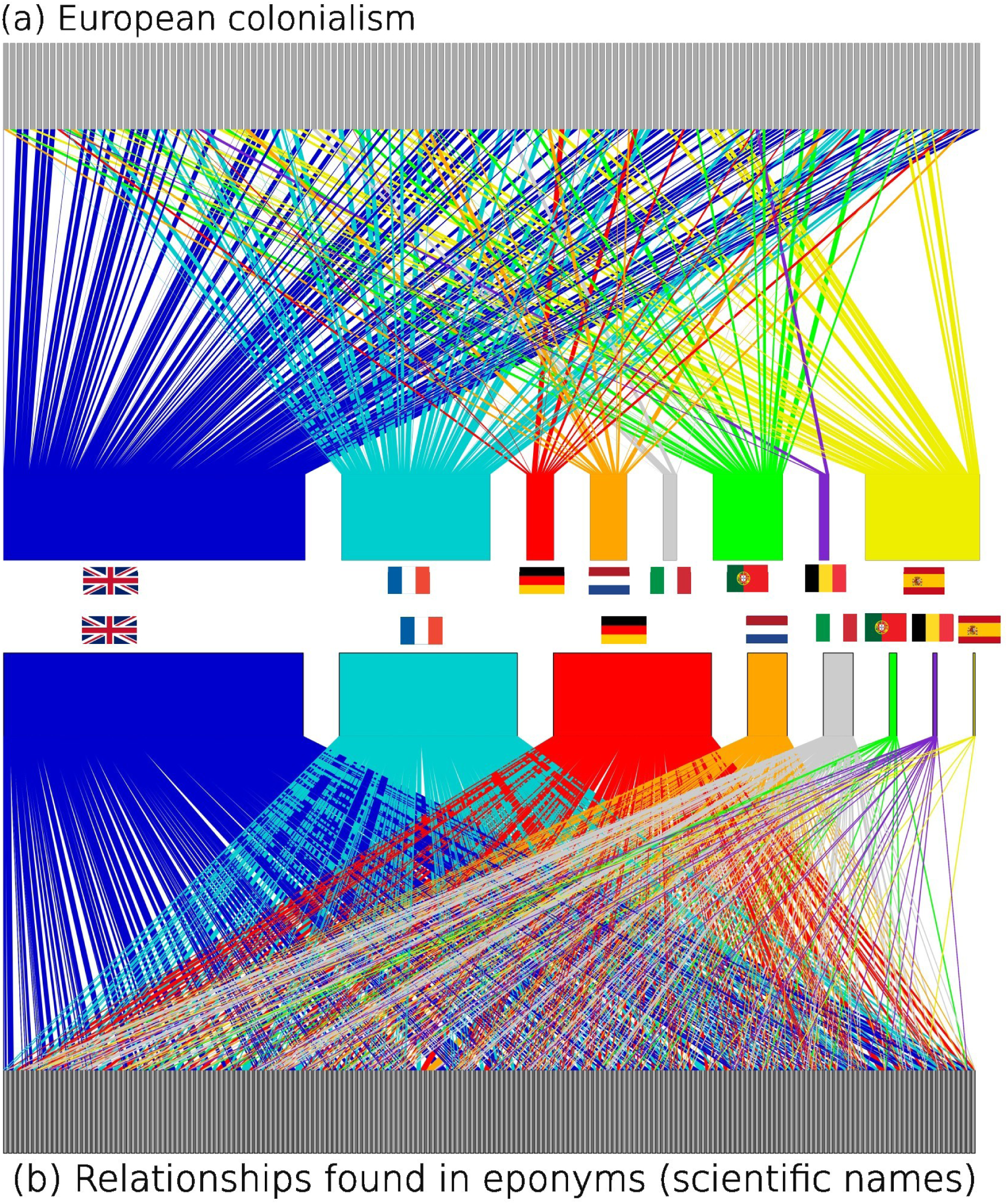
Bipartite network visualization of relationships of (a) historical European colonialism and (b) eponymous bird names. In both networks, the multi-colored layer denotes eight former colonial powers (in order: Blue = Great Britain, Cyan = France, Red = Germany, Orange = Netherlands, Gray = Italy, Green = Portugal, Purple = Belgium, Yellow = Spain), while nodes within other layer – uniformly gray– represents each a modern-day countries (most of which are former colonies, but independent countries today).

**Table 1:** Model estimates of the relationship between national counts of scientific name eponyms and colonizers CEI scores for the country. The model includes observation-level random effects and random intercepts for modern-day countries and colonial empires. R-hat was 1 with effective sample sizes (ESS) over 5000 for all parameters, indicating good convergence and reliable posterior estimation in the model.

| Term | Effect type | Poisson component |  |  | Zero-inflated Component |  |  |
| --- | --- | --- | --- | --- | --- | --- | --- |
|  |  | Estimate | Q2.5 | Q97.5 | Estimate | Q2.5 | Q97.5 |
| Intercept | Fixed | -5.96 | -7.66 | -4.28 | -1.57 | -6.79 | 2.23 |
| CEI | Fixed | 1.33 | 0.47 | 2.30 | 0.32 | -5.32 | 5.64 |
| Colonizer-level intercepts | Random | 2.37 | 1.43 | 4.01 | 19.33 | 6.46 | 51.25 |
| Country-level intercepts | Random | 0.3 | 0.23 | 0.39 | 10.29 | 4.53 | 22.39 |
| Observation-level effect | Random | 0.42 | 0.37 | 0.47 |  |  |  |
| Colonizer-level slopes | Random | 1.23 | 0.67 | 2.21 |  |  |  |
| Correlation between colonizer-specific intercepts and slopes | Random | -0.92 | -1 | -0.61 |  |  |  |

Exampling the effect of CEI among colonizers reveals further interesting details. Persons from countries such as Great Britain and France, were honored in a large number of eponyms overall (higher intercept), even beyond their respective empires (lower slope vis-à-vis CEI). Meanwhile, persons from countries like Spain and Belgium, which were not honored majorly in bird names overall (lower intercepts), were honored predominantly in regions under their respective colonial influences (higher slope vis-à-vis CEI). This effect is further demonstrated by the strong negative correlation between intercepts and slopes (Table 1).

## Discussion

Eponyms are a global phenomenon found in species names across the world, prevalent in the history of global Linnean taxonomy. Of the 2416 eponymous bird species worldwide, a majority honor persons from countries outside the countries where the species occurs (Fig 1a); namely, Great Britain, USA, France and Germany. Overall, a majority of the people honored in eponyms were scientists (40.4%) and naturalists (17.8%), with most eponyms being described in the colonial expansion period. Notably, there is a higher prevalence of country-matching eponyms found in the USA, i.e., American bird names honoring American persons. We also found a shift in the use of eponyms with a trend towards nowadays honoring more native persons from the country where the species is native to.

### Colonial exposure increases bird eponyms attributed to persons of that empire

Greater duration and spatial extents of a specific European colonial occupation (collectively captured by the Colonial Exposure Index [CEI]) increases the number of bird eponyms that honor people from that respective empire (Fig 4). However, this effect varies with the colonial empire – most attributions are towards people from former British and French empires, and far less so for the Spanish and Portuguese empires. Furthermore, eponyms honoring persons from Great Britain, France and Germany were more widespread (i.e., irrespective of their colonial influence in a region), while for the Spanish and Portuguese empires, there was a higher bias towards their own colonies. These patterns are likely associated with the time of their respective expansions. The early phases of European colonialism, dominated by the Spanish and Portuguese colonial powers, focused exclusively on conquest [43]. The description of natural history and scientific knowledge generation only grew in significance towards the late 18^th^ and early 19^th^ century, fueled by British and German ‘explorers’. Throughout the 19th century, European high society developed a growing market for “exotic species”, which was accompanied by the emergence of acclimatization societies to proactively introduce and distribute species across the Empires [44,45]. This institutionalization of colonial trade in biological specimens greatly contributed to increasing the rate of “discovery” of new species, as seen in our results (Fig 2a). Furthermore, technological advancements of the industrial revolution, such as sophisticated guns and steam engine boats, greatly reduced the time and costs of exploration, facilitating an increased rate of specimen collection and species description [46]. This development is represented in bird descriptions like the Prince Ruspoli’s Turraco *Tauraco ruspolii*, shot and collected in southern Ethiopia by the Italian prince, Eugenio Ruspoli.

**Figure 4:**
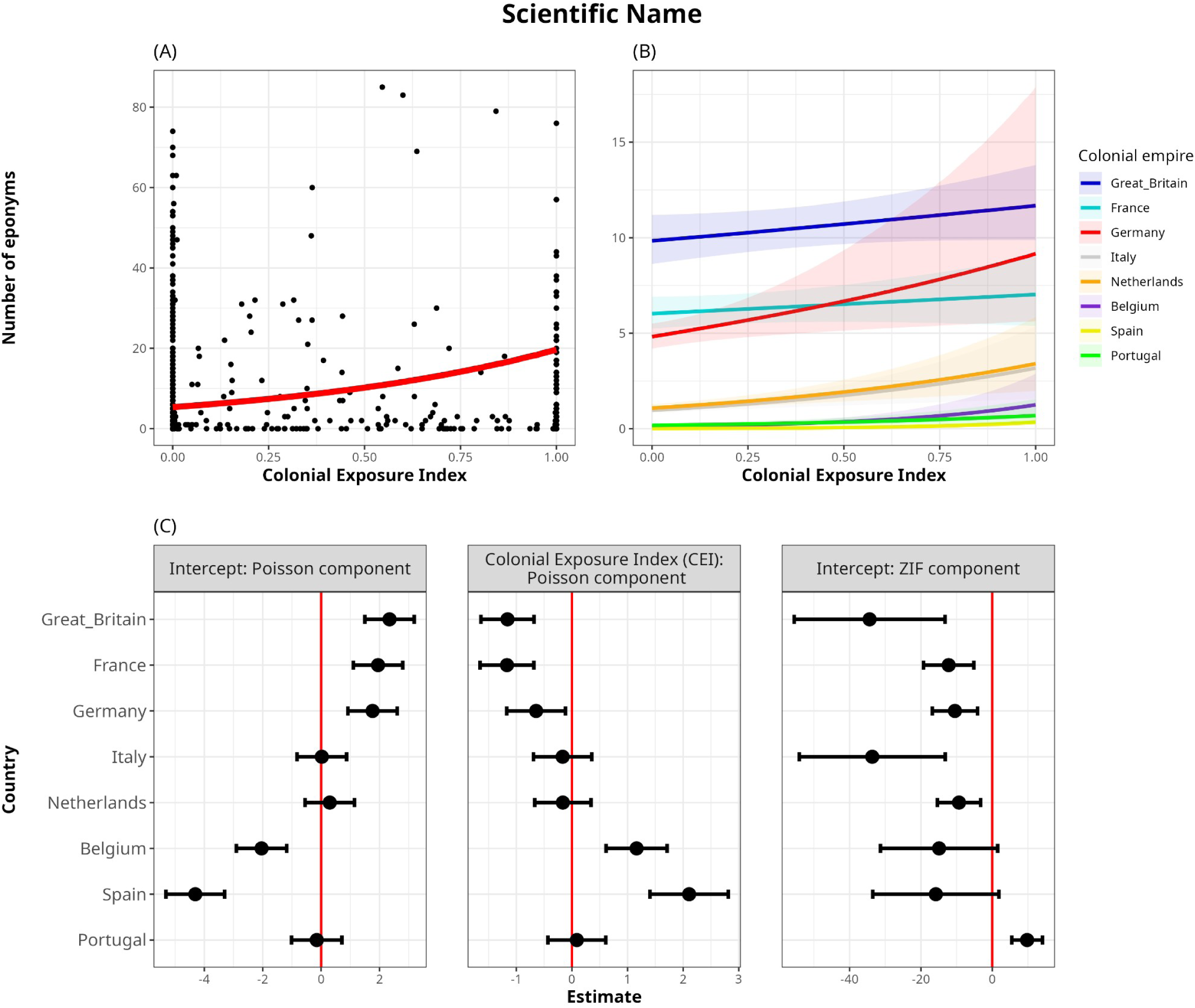
**(A)** Model-predicted relationship between CEI and number of eponyms. Black points represent raw data points, while the red line represents the central tendency of the relationship (posterior median, marginalized over colonizers). There is a four-fold increase in the number of predicted eponyms across the range of CEI values [∼5 eponyms with *CEI = 0*, to ∼20 eponyms with *CEI = 1*]. **(B)** Actual effects of CEI on the number of eponyms (scientific names) for each European colonizer. In the top row, the scale is absolute and represents the actual number of predicted eponyms on the vertical axis. **(C)** Latent scale parameter estimates with the 95% confidence intervals of the Poisson-component intercepts, effect size of Colonial Exposure Index (CEI) and the Zero-inflated-component intercept on the number of eponyms for each colonizer (in scientific names). In the bottom row, the reported values are relative to the overall effect across all colonizers, represented by the posterior-predicted median line shown in (A) and Table 1. Overall, eponyms honoring persons from Great Britain, France and Germany are widespread irrespective of the CEI of the empire in the region; while for the Spanish, Portuguese and Belgian empires, there is a higher number of eponymous bird species honoring its citizens primarily occurring within their own respective former colonies. Statistical support for an overall effect of CEI (i.e. in both components combined) showed a consistent positive effect of CEI on the number of eponyms in >99% of iterations. All model outputs are reported in Table 1.

A high proportion of eponyms globally honor persons from the German empire, despite its smaller colonial extent relative to the British and French empires. Such eponyms honor famous German scientists and naturalists, like the Humboldt’s Hummingbird *Hylocharis humboldtii* honoring Alexander von Humboldt, and Baer’s Pochard *Aythya baeri* honoring Karl Ernst von Baer. Indeed, while eponyms were often associated with peoples of specific occupying colonial powers, influential people in Western science at the time were recognized predominantly in species names. A reason for this could be that the profits of industrialization were channeled towards exploration in certain colonial empires, more so than others [47]. With time, this allowed them to expand their research and exploration efforts across the world, beyond their colonial provinces.

### Shifting regional focus of eponym attribution

Since the 1950s, we see an overall increase in described eponyms honoring native persons and distinguished conservation champions from particular regions. Dugand’s Antwren *Herpsilochmus dugandi* for example, described in 1945, honors the local conservationist Colombian naturalist Armando Dugand. Globally, the diversity of nationalities from which eponyms are selected is also expanding beyond European empires or the USA. Our findings (Fig 2a) at the level of each modern-day country extend those reported by Sangster (2025), who finds that since the 1950s, there is an increasing percentage of bird eponyms honoring people from non-western countries as a whole. That said, many species native ranges transcend political boundaries implying that eponyms will almost inevitably be linked to non-native honorees of some degree.

The USA has the highest number of country-matching eponyms, i.e., cases where the bird species occurs in the same country to which the person honored in its name was from. We speculate that this is due to the USA being a settler society of various European peoples. For example, the Ross’s Goose *Anser rossii* named after an Irish-origin American trader from Boston, Bernard Rogan Ross. Relevant to ongoing debates in the scientific community, this highlights important considerations on when an eponym is to be considered to honor native persons and/or citizens.

### Caveats and important considerations

We acknowledge that assigning quantitative metrics to complex global, historical phenomena such as colonialism and its impacts is an impossible endeavor. We caution that the CEI metric we use is not intended or capable of fully capturing the intricacies and lived realities of the colonial experience. Rather, the CEI is only utilized as a measure for the relative sociopolitical influence of various European empires on each modern-day country, and must only be viewed in relative terms within each country (Fig 5). Further, “country-matching eponyms” highlighted in our study are only defined by the nationality of the person and the countries within the range of the bird species (red edges in Fig 1). Our study does not examine within-country disparities and complexities, notwithstanding that it is possible to have helicopter science operating between distant regions of the same political entity of a country. We also acknowledge that there exist other potentially-related factors that can also influence the number of eponyms in modern-day countries and who they honor, such as migration [48], economic power [14] and science policy [49].

**Figure 5:**
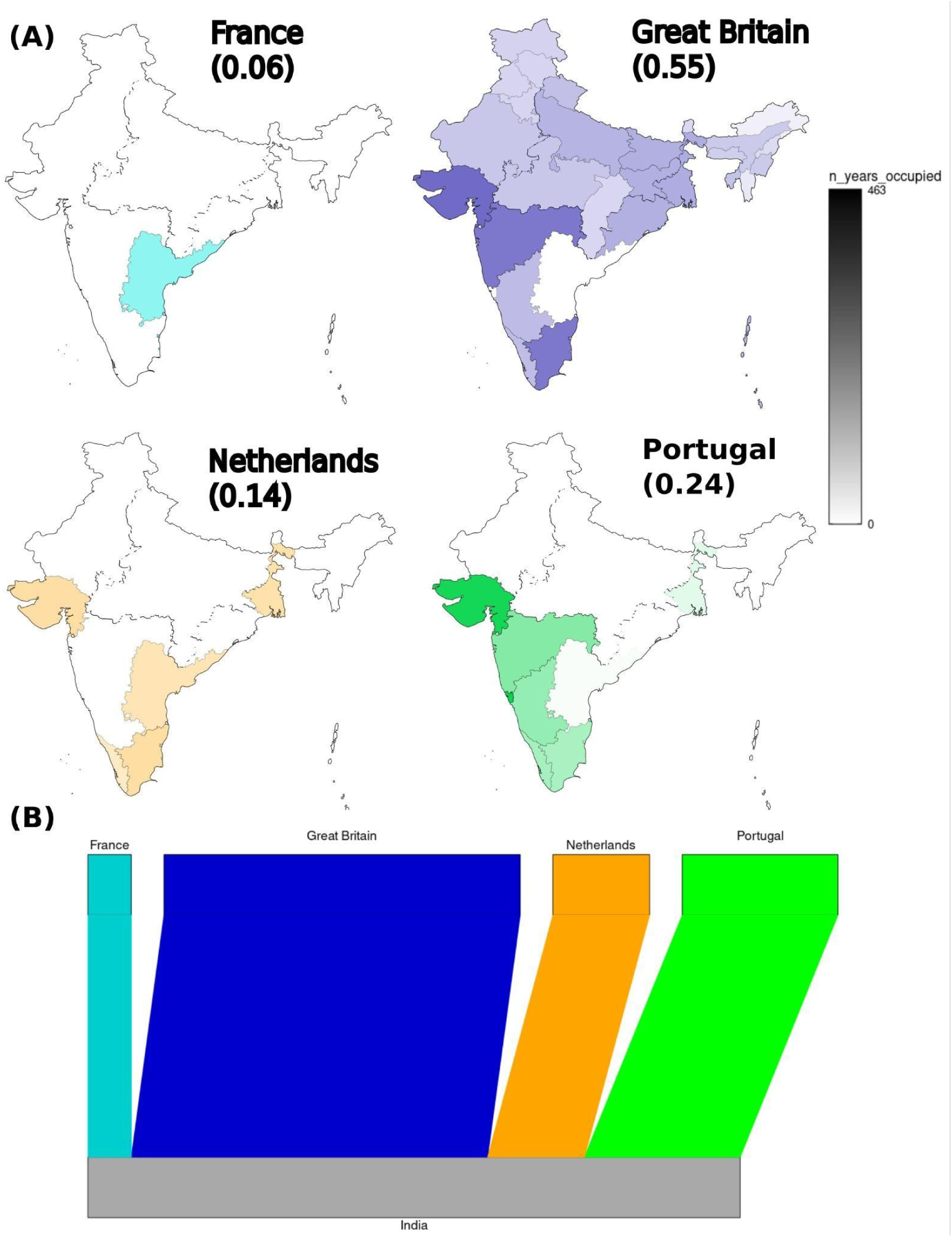
An illustrative example showing how the Colonial Exposure Index of various European empires was calculated for India, a country which was colonized by multiple European empires during the colonial times. **(A)** Each empire is shown in a separate panel indicated by a different color: France (cyan), Great Britain (blue), Netherlands (orange) and Portugal (green). The intensity of the color represents the number of years that the colonial power was influential or in control of the region; **(B)** A bipartite network representation of the colonialism experienced by India by European colonial powers; the width of each edge represents the CEI associated with its colonization of India.

Eponymous migratory species present an important consideration when discussing the future of eponym usage worldwide. Each year, migratory species spend their time across their breeding, migration and wintering grounds across vast geographic ranges, and in the process bridge distant countries and their historical legacies. Consequently, the presence of migratory large-ranging species leads to increased occurrence of ‘non-native’ eponym usage, simply by virtue of being present (at some point in the year) in various modern-day countries. For example, the Tickell’s Leaf Warbler *Phylloscopus affinis,* a bird named after British colonel Samuel Richard Tickell, breeds in the Himalayas and central-China and winters across the Indian subcontinent in India, Bangladesh, Myanmar and Thailand. Its range encompasses countries and regions with vastly different historical contexts and cultures.

Migratory species [nearly 17.9% of all extant bird species; 1997 species, 50] are not a discrepancy, but highlight that taxonomic decisions are made at the global level, across the entire geographic range of each species. It brings into question elements of the Guedes et al. (2023) proposal to eradicate eponyms altogether. Notably, there are relatively fewer such eponymous migratory species that are ‘country-matching’ when found in formerly-colonizer countries (notice the near-absence of red lines in Fig 1a originating from formerly-colonizer countries). Indeed, a considerable fraction of European names predate the widespread acceptance of eponyms in bird nomenclature. Nevertheless, to understand the statistical importance of migratory species in the eponym-CEI relationship, we performed a sensitivity analysis after excluding all migratory species found in the main dataset [50]. We find no significant differences between interpretations drawn from this analysis and our main model, highlighting that the trends seen in the eponym-CEI relationship are not biased by the inclusion of migratory species (Supplementary Analysis 2: Fig S7; Table S8).

Overall, our study provides the first network-based, statistical examination of bird eponyms worldwide and their colonial legacies, which was hitherto lacking in lieu of recent debates that examine etymologies in a case-to-case and/or prescriptive manner. Our findings extend those of Du Bay et al. (2020) and Sangster (2025) who analyze patterns in global bird eponym usage (in scientific names). Our network-based approach enables understanding more nuanced relationships between the European colonial legacy of a country and its bird eponyms today at finer national and sub-national spatial scales.

## Methods

### Bird eponyms

The list of all bird names and their distributions was accessed from BirdLife International HBW Version 8.1 [50], a widely used resource for species names and corresponding distributions of bird species. Of the total 11,195 bird species listed, we analyzed only the species with available range maps for their distributions, i.e., 11,068 species. Previous versions of the list (likely with more eponymous bird names in them) were not analyzed. Maps of subnational regions and national boundaries follow the Biodiversity Information Standards database of Global Administrative Areas (GADM) for country level entities and the geographical scheme level 4 for subnational entities [51].

By overlapping the range maps of each species with the national boundaries, a national list of all bird species found within each country was collated. For our analyses, we excluded all non-native ranges of species and regions with vagrant reports of the species. Migratory species were listed in each national list across their migratory range. All data filtering steps were performed using the *tidyverse* package [Version 2.0.0, 52] in the R programming language [53]. Spatial data was processed using the *sf* package [Version 1.0.21, 54].

All 2416 bird eponyms, across the common and the scientific names, were marked through thorough manual inspection of *The eponym dictionary of birds* [55] and subsequent online searches. Additionally, from the text presented in the book, the name, nationality and profession of the person honored in each eponym was noted, alongside the year of species description. In many cases, the professions encompassed several disciplines and domains, for example, Charles Joseph Bernier (honored in Bernier’s Vanga *Oriolia bernieri*) was a surgeon, botanist and collector. In such cases, a hierarchy of descriptions (listed in Table S5) was utilized to mark the profession for each person honored in the eponym.

We assigned the nationality of honored persons by following the decision tree in Fig S2. The decision tree helps to reproducibly assign and transparently report our methodology, particularly regarding persons who migrated during their lives – such as that of Bernard Rogan Ross, an Irish-born trader who spent his life working and living in modern-day Canada. Furthermore, we examined the sensitivity of our results and interpretations to the decisions made when assigning nationality by birth vis-à-vis using our decision tree (Supplementary Analysis 1). We find no major differences between results interpretations from analyses run on both approaches (Supplementary Analysis 1: Fig S6; Table S7). Of the 2416 eponyms we analyzed, only 4.5% were assigned different nationalities of honored persons when using nationality-by-birth or using our decision tree. The full list of the analyzed eponyms, the nationality of the honored person, year of description and their profession/relationship with describer can be found in the Supplementary Materials.

### European colonialism

Data on European colonialism was accessed from Lenzner et al. (2022). This dataset contains information on the colonial history of most “regions” on earth, each defined as a (sub)national identity which was under a particular colonial rule. In some cases in the absence of more fine scale data, the whole area of a modern day country (Fig S1) was demarcated as a single region; for example, the country of Nigeria. However, in many cases, a region encompasses a smaller sub-national region which has a shared colonial legacy, for example, the states of modern-day USA and India. To demarcate national territories, we combined all regions found within a single country into a single entity. Regions where the whole country was demarcated as a singular region remain unchanged in this merging step (Fig S1).

Subsequently, we calculated a **Colonial Exposure Index (CEI)** for each colonizer in each formerly-colonized country (as defined by modern boundaries). The CEI of a particular colonizer *j* on a particular modern-day country *k* (with *n* regions contained within its borders today) is defined as follows:

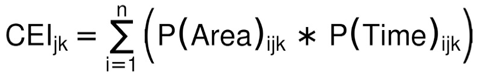

where,

P(Area)*_ijk_*= proportion of area occupied by region *i* relative to the total area of the country (occupied by colonizer *j* in modern-day country *k*)
P(Time)*_ijk_* = proportion of time that region *i* was occupied by colonizer *j* in modern-day country *k*, relative to the total time period the country *k* experienced any colonial rule

The CEI metric factors in both the duration of occupation that a particular European empire had in a country and the proportion of its total area which it occupied. For example, India experienced colonization by several European colonial powers (British, French, Portuguese and Dutch colonization; Fig 5a). The CEI of India under British colonization is 0.55, the highest among all European empires (since the British empire occupied most of modern day India). The CEI of Portuguese colonization in India is also noticeable (CEI = 0.24) despite the fact that the Portuguese occupied a much smaller territory. This is because certain territories were occupied for a long period of time, e.g., 451 years in the case of Goa.

The CEI metric does not calibrate with calendar dates, and is agnostic to trends in taxonomic nomenclature or other social events. It collates the information on area and time of a specific colonial influence since early European exploration to the post-colonial independence of modern-day countries. The principle utility of CEI is to estimate the relative influence of each former empire on each modern-day country. Since CEI is designed as a predictor variable that can be calculated across context for every modern-day country, it involves a date-agnostic calculation. Statistically, this ensures that there is no forced correlation by virtue of the definition of CEI, between our predictor (measures of colonial influence) and response variables (number of bird eponyms). This definition of CEI also makes it useful for investigating other modern-day phenomena stemming from region-specific colonial histories.

### Bipartite network analyses

Our primary objective was to explicitly test the effect of European colonization on the world’s bird eponyms today. We use network analyses owing to its compatibility with our research questions, i.e., since the identities of countries (former colonizer and colonized countries) was maintained in the colonialism and eponym datasets. Furthermore, quantitative techniques to test the effect of one network on another are well-described from other ecological studies [56].

In both scientific and common names, we constructed a weighted bipartite network using the *igraph* package [Version 2.1.4, 57] in the R programming language [Version 4.5.1, 53]. Common names and scientific names were analyzed separately throughout. The relationships were visually examined in two forms: (a) relationships across all countries (including eponyms honoring persons from non-colonial countries); (b) visualization of relationships only among eponyms honoring persons from European colonial powers.

We further analyzed the latter relationship statistically using the *igraph*, *bipartite* [Version 2.21, 58] and *brms* packages [Version 2.23.0, 59]. First, we constructed a bipartite network for European colonialism – where the nodes represented colonizer and colonized countries, and each weighted edge between them represents the CEI of the specific colonizer on the specific colonized country (Fig 5). Separately, we constructed a separate bipartite network for relationships found in bird eponyms, where the nodes were the ‘colonizer’ countries (here, country to which the persons honored in the eponym belonged to) and the ‘colonized’ countries (here, countries where the eponymously-named bird species occurs). Edge weights between nodes represent the number of eponyms honoring a person from the colonizer country which is found in that colonized country.

Using these networks with comparable structure, we model the colonialism network as a predictor of the number of eponyms (response). Specifically, we use Bayesian generalized linear mixed models in the *brms* package [Version 2.23.0, 59]. Due to a larger number of zeros than expected from an overdispersed poisson, we modeled this as a zero-inflated poisson, with a log link function on the poisson component and a logit link on the zero-inflated (binomial) component (see supplementary materials). We modeled a fixed effect of CEI and random country and colonizer effects for both poisson and zero-inflated components. For the poisson component, we additionally modeled random slopes of CEI for every colonizer, which allows the CEI to vary across colonizers, and included an offset term to account for different countries having different numbers of species in their national lists (and consequently, the probability of describing a species varying across countries). We also included an observation level random effect to account for overdispersion [60]. We use weak, uninformative priors as implemented in *brms*. Fixed effects were assigned normal priors with mean 0 and standard deviation 10. Random intercepts and slope standard deviations were assigned Student-t priors with 3 degrees of freedom, mean 0, and scale 10. Correlations among group-level effects were assigned LKJ(1) priors. Offsets were treated as fixed constants. Four chains were run for each model, with a warmup of 2000 iterations and 10000 iterations post warmup. Model parameters were assessed based on the Gelman-Rubin diagnostic [R^W^; 61], ESS values and diagnostic plots (reported in supplementary material) made using the *ggplot2* package [Version 3.5.2, 52]. In order to test the statistical support for the total relationship between CEI and the number of eponyms, we predicted the number of eponyms on the observed scale across the range of CEI (0-1; see Figure 4a) and then calculated the proportion of iterations in which the relationship (slope) was positive (see Supplementary Methods for detailed description).

#### Supplementary analyses 1: Assigning nationality by birth (not following the decision tree)

To understand the sensitivity of our analyses to decisions made using our decision tree (Fig S2), we separately re-analyzed our study after assigning nationality based on the country of birth – irrespective of the reason of being in the country or if they became naturalized citizens. Our study uses information on persons, as provided in the *The eponym dictionary of birds* [55], and occasionally on AviList [62]. Overall, of the 2416 eponyms we analyzed, only 108 (4.5%) were assigned different nationalities of honored persons when using nationality-by-birth vs. using the decision tree. The model outputs and predicted relationships are shown in Fig S6 and Table S7.

#### Supplementary analyses 2: Excluding migratory species

To understand the sensitivity of our analyses to migratory species presence across different countries, separately re-analyzed our study after excluding all migratory species and their ranges, as mentioned in BirdLife International HBW [50]. We performed the same analytical workflows and statistical reporting as in our main analysis. The model outputs and predicted relationships are shown in Fig S7 and Table S8.

## Acknowledgements

We thank Dr Matt Silk for the initial discussions on network analyses, and Dr Anand Krishnan for the comments on an earlier version of the manuscript.

## Funding information

We did not receive any specific funding for this work. Open access publishing was facilitated by Universite de Neuchatel, via the Consortium Of Swiss Academic Libraries. Akshay Bharadwaj is grateful for funding from the Swiss National Science Foundation (SNSF). Bernd Lenzer appreciates funding from Austrian Science Fund FWF (Global Plant Invasions, grant no. I 5825- B).

## Data sharing statement

All scripts, shapefiles and datasets necessary to fully reproduce our figures and results will be published with a DOI on Zenodo.

## Author contributions

Akshay Bharadwaj ideated and designed the study. Bernd Lenzer contributed the colonialism datasets and shapefiles he had compiled and curated. Akshay manually annotated eponym information, and performed the statistical analyses with guidance from Joel Pick. Akshay Bharadwaj wrote the first draft of the manuscript, while Bernd Lenzner and Joel Pick aided in its revision.

## Competing interests statements

The authors declare no competing interests.

## Notes

### Competing Interest Statement

The authors have declared no competing interest.

## References

1. Roberts L. Situating Science in Global History: Local Exchanges and Networks of Circulation. Itinerario. 2009;33: 9–30. doi:10.1017/S0165115300002680

2. Seth S. Putting knowledge in its place: science, colonialism, and the postcolonial. Postcolonial Studies. 2009;12: 373–388. doi:10.1080/13688790903350633

3. Engerman SL, Sokoloff KL. Colonialism, inequality, and long-run paths of development. National Bureau of Economic Research Cambridge, Mass., USA; 2005.

4. Levy JT. Colonialism and Its Legacies. Lexington Books; 2011.

5. Baber Z. The Plants of Empire: Botanic Gardens, Colonial Power and Botanical Knowledge. Journal of Contemporary Asia. 2016;46: 659–679. doi:10.1080/00472336.2016.1185796

6. Braby MF, Hsu Y-F, Lamas G. How to describe a new species in zoology and avoid mistakes. Zool J Linn Soc. 2024;202: zlae043. doi:10.1093/zoolinnean/zlae043

7. Green M. Gone Cuckoo. Biodiversity. 2018;19: 216–218. doi:10.1080/14888386.2018.1512419

8. Collar N, Christie D, Goodman DD, Pyle P, Kirwan GM, Boesman PFD. Common Nightingale (Luscinia megarhynchos), version 1.1. Birds of the World. 2025 [cited 27 Mar 2026]. doi:10.2173/bow.comnig1.01.1

9. He L, Cornish TC, Kricka LJ, Vandergriff TW, Yancey K, Nguyen K, et al. Trends in dermatology eponyms. JAAD Int. 2022;7: 137–143. doi:10.1016/j.jdin.2022.03.006

10. Armocida E, Masciangelo G, Natale G. Medical eponyms versus acronyms: what medical terminology is most beneficial to learn? A question of goals. Postgraduate Medical Journal. 2024;100: 771–775.

11. Manz N, McCullough I. Eponyms in Science: how long can they get? Scientometrics. 2025;130: 3455–3482. doi:10.1007/s11192-025-05328-9

12. DuBay S, Palmer DH, Piland N. Global inequity in scientific names and who they honor. BioRxiv. 2020; 2020.08. 09.243238.

13. Chen-Kraus C, Farmer C, Guevara EE, Meier K, Watts DP, Widness J. Whom Do Primate Names Honor? Rethinking Primate Eponyms. Int J Primatol. 2021;42: 980– 986. doi:10.1007/s10764-021-00252-0

14. Abreu ECT, da Silva EL, Moura MR. Geopolitical impacts on the description of new terrestrial mollusc species. Proc Biol Sci. 2025;292: 20251428. doi:10.1098/rspb.2025.1428

15. Beolens B, Watkins M, Grayson M. The eponym dictionary of amphibians. Pelagic Publishing; 2013.

16. Beolens B, Watkins M, Grayson M. The eponym dictionary of reptiles. JHU Press; 2011.

17. Ceríaco LMP, Aescht E, Ahyong ST, Ballerio A, Bouchard P, Bourgoin T, et al. Renaming taxa on ethical grounds threatens nomenclatural stability and scientific communication: Communication from the International Commission on Zoological Nomenclature. Zoological Journal of the Linnean Society. 2023;197: 283–286. doi:10.1093/zoolinnean/zlac107

18. Antonelli A, Farooq H, Colli-Silva M, Araújo JPM, Freitas AVL, Gardner EM, et al. People-inspired names remain valuable. Nat Ecol Evol. 2023;7: 1161–1162. doi:10.1038/s41559-023-02108-7

19. Guedes P, Alves-Martins F, Arribas JM, Chatterjee S, Santos AMC, Lewin A, et al. Eponyms have no place in 21st-century biological nomenclature. Nat Ecol Evol. 2023;7: 1157–1160. doi:10.1038/s41559-023-02022-y

20. Jost L, Yanez-Muñoz MH, Brito J, Reyes-Puig C, Reyes-Puig JP, Guayasamín JM, et al. Eponyms are important tools for biologists in the Global South. Nat Ecol Evol. 2023;7: 1164–1165. doi:10.1038/s41559-023-02102-z

21. Mabele MB, Kiwango WA, Mwanyoka I. Disrupting the epistemic empire is necessary for a decolonial ecology. Nat Ecol Evol. 2023;7: 1163–1163. doi:10.1038/s41559-023-02105-w

22. Orr MC, Hughes AC, Carvajal OT, Ferrari RR, Luo A, Rajaei H, et al. Inclusive and productive ways forward needed for species-naming conventions. Nat Ecol Evol. 2023;7: 1168–1169. doi:10.1038/s41559-023-02103-y

23. Roksandic M, Musiba C, Radović P, Lindal J, Wu X-J, Figueiredo E, et al. Change in biological nomenclature is overdue and possible. Nat Ecol Evol. 2023;7: 1166– 1167. doi:10.1038/s41559-023-02104-x

24. Thiele K. Some, but not all, eponyms should be disallowed. Nat Ecol Evol. 2023;7: 1170–1170. doi:10.1038/s41559-023-02106-9

25. Bae CJ, Radović P, Wu X-J, Figueiredo E, Smith GF, Roksandic M. Placing taxonomic nomenclatural stability above ethical concerns ignores societal norms. Zoological Journal of the Linnean Society. 2023;199: 5–6. doi:10.1093/zoolinnean/zlad061

26. American Ornithological Society. American Ornithological Society Will Change the English Names of Bird Species Named After People. In: American Ornithological Society [Internet]. 1 Nov 2023 [cited 29 Jan 2025]. Available: https://americanornithology.org/american-ornithological-society-will-change-the-english-names-of-bird-species-named-after-people/

27. Sangster G. Eponyms of birds mostly honour scientists and show positive inclusivity trends. Zoological Journal of the Linnean Society. 2025;203: zlaf022. doi:10.1093/zoolinnean/zlaf022

28. Liu IA, Gulson-Castillo ER, Wu JX, Demery A-JC, Cortes-Rodriguez N, Covino KM, et al. Building bridges in the conversation on eponymous common names of North American birds. Ibis. 2024;166: 1092–1102. doi:10.1111/ibi.13320

29. Stefanoudis PV, Licuanan WY, Morrison TH, Talma S, Veitayaki J, Woodall LC. Turning the tide of parachute science. Current Biology. 2021;31: R184–R185. doi:10.1016/j.cub.2021.01.029

30. Asase A, Mzumara-Gawa TI, Owino JO, Peterson AT, Saupe E. Replacing “parachute science” with “global science” in ecology and conservation biology. Conservation Science and Practice. 2022;4: e517. doi:10.1111/csp2.517

31. Groom Q, Meeus S, Bárrios S, Childs C, Clubbe C, Corbett E, et al. Capacity building needed to reap the benefits of access to biodiversity collections. PLANTS, PEOPLE, PLANET. 2025;n/a. doi:10.1002/ppp3.70029

32. Raposo MA, da Silva HR, Francisco BCS, Vieira O, da Fonseca OV, de Assis CP, et al. Is stability too revered in zoological nomenclature? Zoological Journal of the Linnean Society. 2023;199: 7–9. doi:10.1093/zoolinnean/zlad106

33. Pethiyagoda R. Policing the scientific lexicon: The new colonialism?. Megataxa. 2023;10: 20–25. doi:10.11646/megataxa.10.1.4

34. Jiménez-Mejías P, Manzano S, Gowda V, Krell F-T, Lin M-Y, Martín-Bravo S, et al. Protecting stable biological nomenclatural systems enables universal communication: A collective international appeal. BioScience. 2024;74: 467–472. doi:10.1093/biosci/biae043

35. He Z-Q. Let scientific names and indigenous names carry out their respective duties. Zoological Journal of the Linnean Society. 2025;203: zlae063. doi:10.1093/zoolinnean/zlae063

36. Smith GF, Figueiredo E. “Rhodes-” must fall: Some of the consequences of colonialism for botany and plant nomenclature. TAXON. 2022;71: 1–5. doi:10.1002/tax.12598

37. Bascompte J. Networks in ecology. Basic and Applied Ecology. 2007;8: 485–490. doi:10.1016/j.baae.2007.06.003

38. Borrett SR, Moody J, Edelmann A. The rise of Network Ecology: Maps of the topic diversity and scientific collaboration. Ecological Modelling. 2014;293: 111–127. doi:10.1016/j.ecolmodel.2014.02.019

39. Giannini TC, Garibaldi LA, Acosta AL, Silva JS, Maia KP, Saraiva AM, et al. Native and non-native supergeneralist bee species have different effects on plant-bee networks. PloS one. 2015;10: e0137198.

40. Ride W. International code of zoological nomenclature. International Trust for Zoological Nomenclature; 1999.

41. Driver RJ, Bond AL. Towards redressing inaccurate, offensive and inappropriate common bird names. Ibis. 2021;163: 1492–1499. doi:10.1111/ibi.12984

42. Andrade R, Larson KL, Franklin J, Lerman SB, Bateman HL, Warren PS. Species traits explain public perceptions of human–bird interactions. Ecological Applications. 2022;32: e2676. doi:10.1002/eap.2676

43. Wood L. The environmental impacts of colonialism. 2015.

44. Anderson W. Climates of opinion: acclimatization in nineteenth-century France and England. Victorian studies. 1992;35: 135–157.

45. Osborne MA. Acclimatizing the world: a history of the paradigmatic colonial science. Osiris. 2000;15: 135–151.

46. Johnson K. Type-specimens of birds as sources for the history of ornithology. Journal of the History of Collections. 2005;17: 173–188. doi:10.1093/jhc/fhi027

47. Stafford RA. Scientific Exploration and Empire. The Rise and Fall of Modern Empires, Volume II. Routledge; 2013.

48. Niva V, Horton A, Virkki V, Heino M, Kosonen M, Kallio M, et al. World’s human migration patterns in 2000–2019 unveiled by high-resolution data. Nature Human Behaviour. 2023;7: 2023–2037.

49. Taylor MZ. The politics of innovation: Why some countries are better than others at science and technology. Oxford University Press; 2016.

50. HBW and BirdLife International. Handbook of the Birds of the World and BirdLife International digital checklist of the birds of the world. Version 8.1. 2024 [cited 29 Jan 2025]. Available: Available at http://datazone.birdlife.org/species/requestdis.

51. Brummitt RK, Pando F, Hollis S, Brummitt NA. World geographical scheme for recording plant distributions. International working group on taxonomic databases for plant sciences; 2001.

52. Wickham H, Averick M, Bryan J, Chang W, McGowan LD, François R, et al. Welcome to the Tidyverse. Journal of open source software. 2019;4: 1686.

53. R Development Core Team. R: A language and environment for statistical computing. 2023.

54. Pebesma E, Bivand R. Spatial data science: With applications in R. Chapman and Hall/CRC; 2023.

55. Beolens B, Watkins M, Grayson M. The eponym dictionary of birds. Bloomsbury Publishing; 2020.

56. Juárez-Juárez B, Dáttilo W, Moreno CE. Synthesis and perspectives on the study of ant-plant interaction networks: A global overview. Ecological Entomology. 2023;48: 269– 283. doi:10.1111/een.13227

57. Csardi G, Nepusz T. The igraph software package for complex network research. InterJournal, complex systems. 2006;1695: 1–9.

58. Dormann CF, Fründ J, Blüthgen N, Gruber B. Indices, graphs and null models: analyzing bipartite ecological networks. 2009.

59. Bürkner P-C. brms: An R Package for Bayesian Multilevel Models Using Stan. Journal of Statistical Software. 2017;80: 1–28. doi:10.18637/jss.v080.i01

60. Hinde J. Compound Poisson regression models. Glim 82: Proceedings of the international conference on generalised linear models. Springer; 1982. pp. 109–121.

61. Gelman A, Rubin DB. Inference from iterative simulation using multiple sequences. Statistical science. 1992;7: 457–472.

62. AviList Core Team. AviList: The Global Avian Checklist, v2025. 2025. Available: 10.2173/avilist.v2025

