## Supplementary File for "Colonial legacies in eponymous species names: a global network perspective"

**Supplementary materials:**

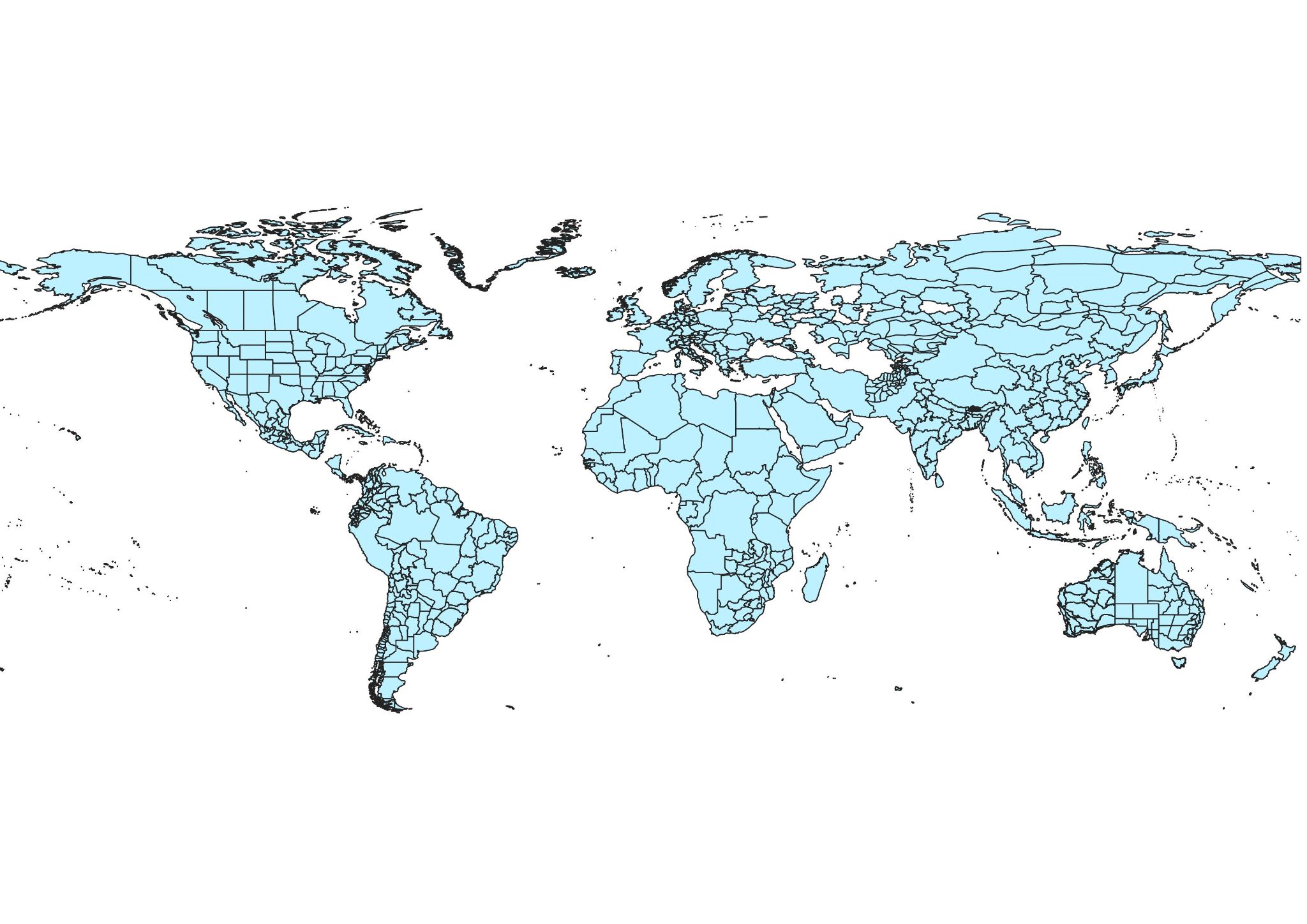
**Figure S1: A global map showing the analyzed regions. Some countries (such as Nigeria) encompass the area of the entire country. Meanwhile, other countries (such as India), consist of multiple “regions” which each represent an area with a shared colonial history.**

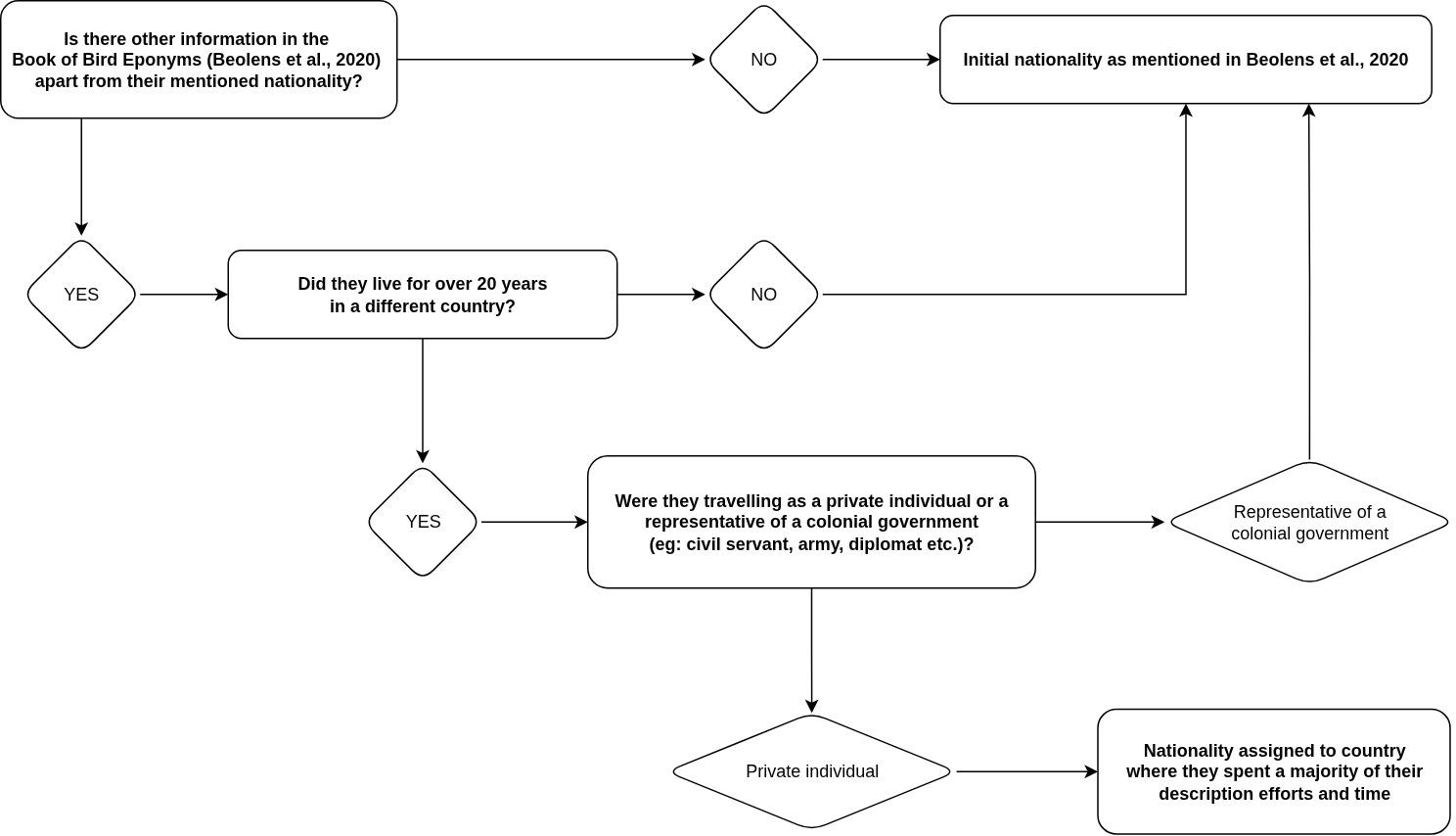

**Figure S2:** The decision tree followed to assign nationality of honored persons. Nationality (Beolens et al., 2020) was updated only when a person from a certain country (often a colonial empire) spent over two decades living and working in foreign country in a private capacity, i.e., not part of a colonial administrative service (diplomacy, civil service, military tenure etc.). We utilized the decision tree to help assign nationalities in a transparent and reproducible manner.

*Citation:*

Beolens, B., Watkins, M., & Grayson, M. (2020). *The eponym dictionary of birds*. Bloomsbury Publishing.

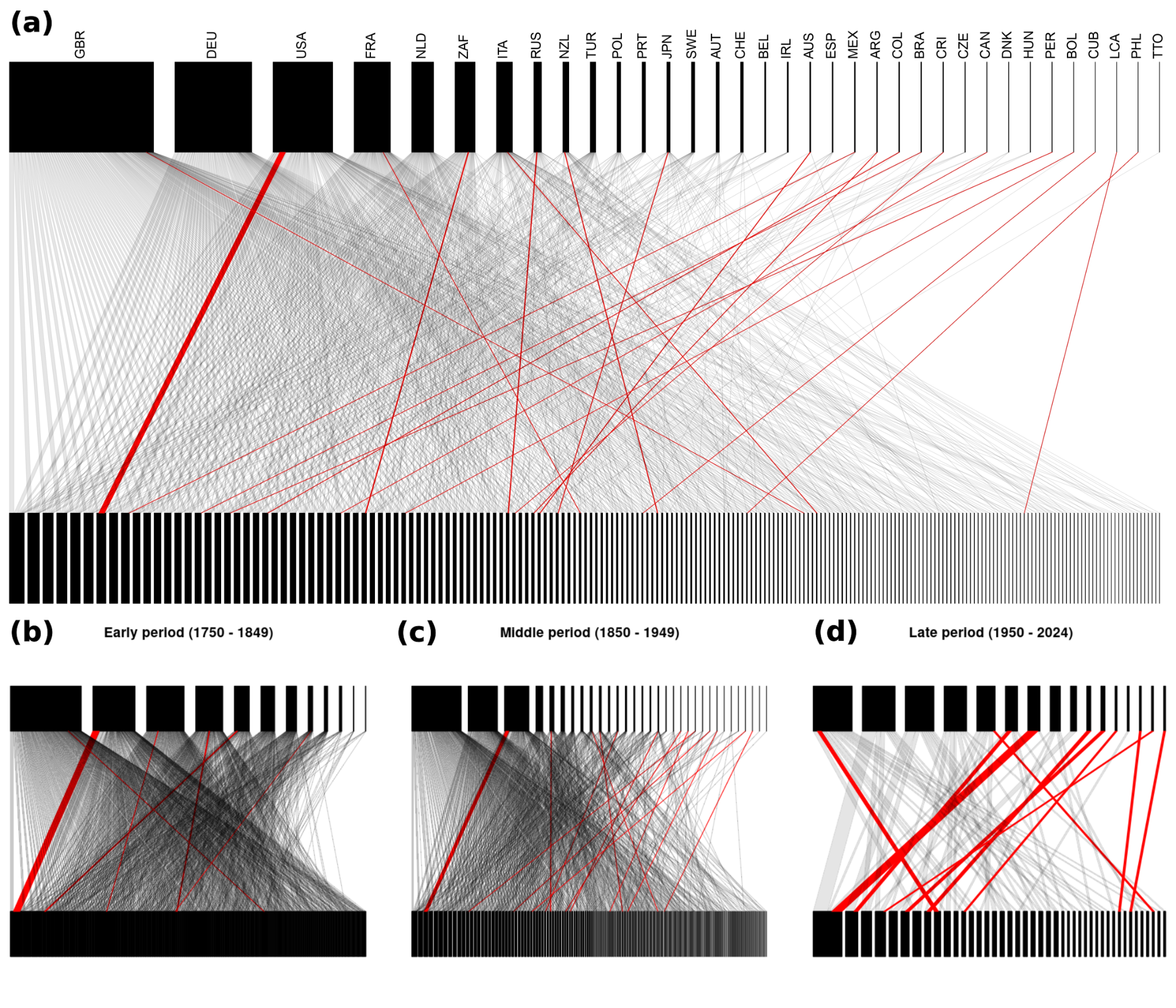

**Figure S3:** Bipartite networks visualizing relationships in eponymous common bird names (a) overall, (b) in the early colonization period, (c) colonial expansion period and (d) post-colonial period. In each separate network, countries where eponymous birds occur are shown in the bottom layer, while the country of origin of the persons honored in the eponymous bird name is shown on the top layer. A gray edge between the layers denotes a single eponymous bird species found in a country when it is named after a person who is not from that country. Conversely, red edges highlight cases where the bird and the person honored in its name are from the same country. *The full key to all ISO country codes is provided in Table S1.*

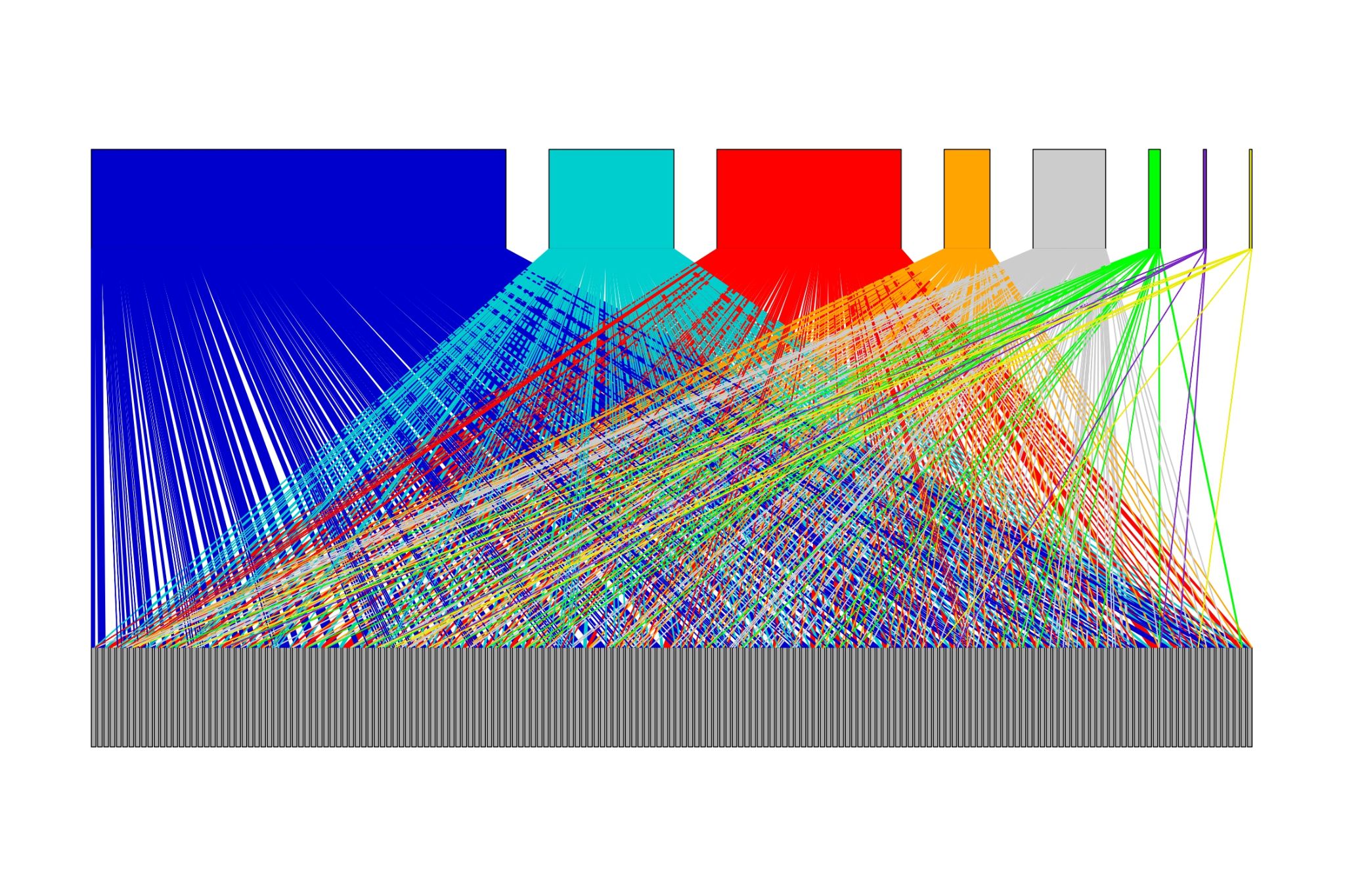

**Figure S4:** Bipartite network visualization of relationships of eponymous bird names (common names in English). The top layer denote former colonial powers *(in order: Blue = Great Britain, Cyan = France, Red = Germany, Orange = Netherlands, Gray = Italy, Green = Portugal, Purple = Belgium, Yellow = Spain*), while gray nodes in the bottom layer represent former colonies which are today independent political entities.

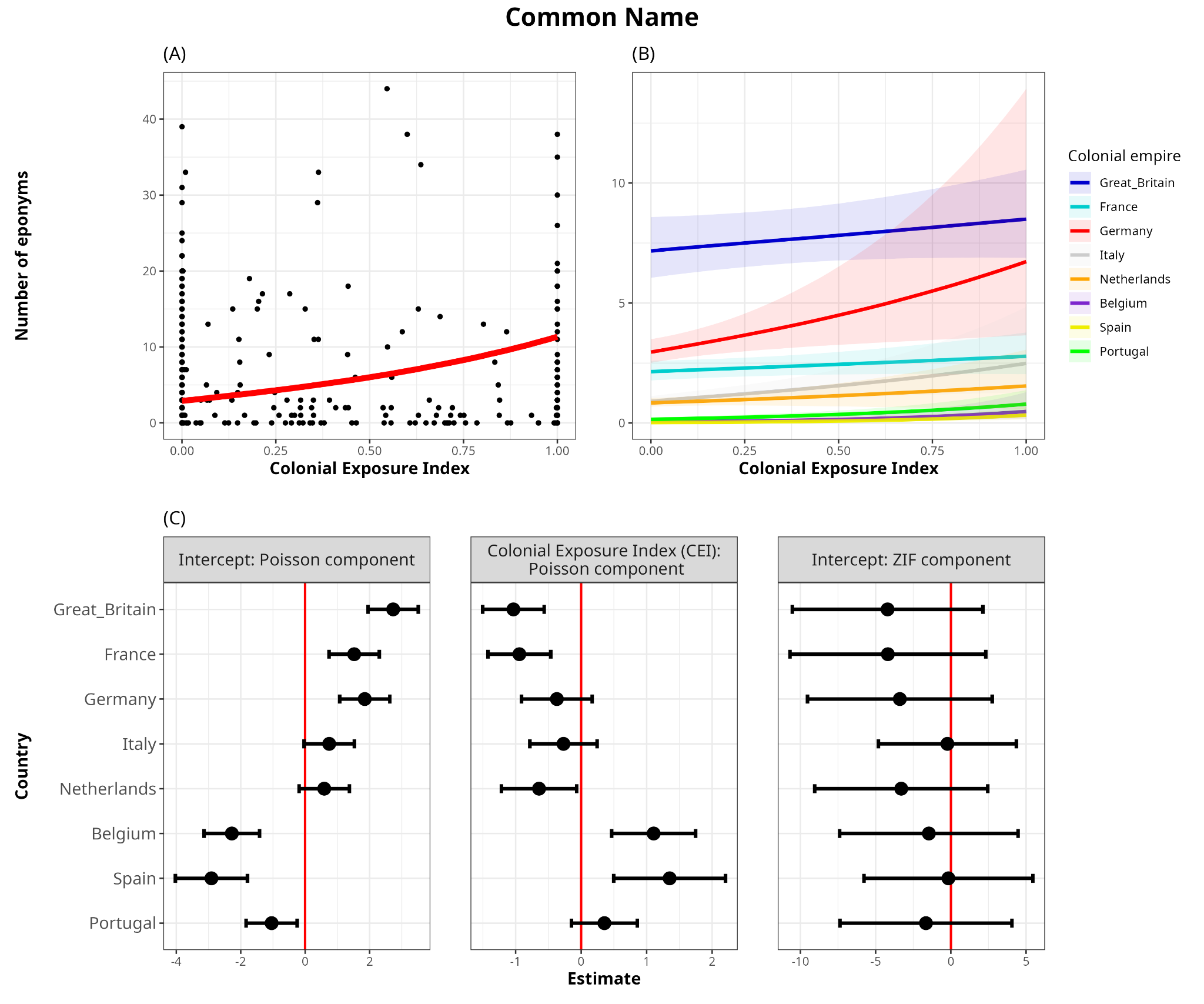

**Figure S5:** **(A)** Model-predicted relationship between CEI and number of eponyms in common names. Black points represent raw data points, while the red line represents the central tendency of the relationship (posterior median, marginalized over colonizers). There is a three-fold increase in the number of predicted eponyms across the range of CEI values [from ~3 eponyms with *CEI = 0*, to 11 eponyms with *CEI = 1*]. **(B)** Actual effects of CEI on the number of eponyms (common names) for each European colonizer. In the top row, the scale is absolute and represents the actual number of predicted eponyms on the vertical axis. **(C)** Latent scale parameter estimates with the 95% confidence intervals of the Poisson-component intercepts, effect size of Colonial Exposure Index (CEI) and the Zero-inflated-component intercept on the number of eponyms for each colonizer (in common names). In the bottom row, the reported values are relative to the overall effect across all colonizers, represented by the posterior-predicted median line shown in (A) and Table 1. Overall, eponyms honoring persons from Great Britain, France and Germany are widespread irrespective of the CEI of the empire in the region; while for the Spanish, Portuguese and Belgian empires, there is a higher number of eponymous bird species honoring its citizens primarily occurring within their own respective former colonies. Statistical support for an overall effect of CEI (i.e. in both components combined) showed a consistent positive effect of CEI on the number of eponyms in >99% of iterations. All model outputs are reported in Table S6.

**Table S1: Number of eponyms and the number of eponyms as a percentage of total national bird richness of each country.**

| **3-letter ISO code** | **Country** | **Total number of bird species** | **Number of eponyms in scientific name** | **Percentage of country's bird scientific names that are eponymous** | **Number of eponyms in common name** | **Percentage of country's bird common names that are eponymous** |
| --- | --- | --- | --- | --- | --- | --- |
| **ABW** | **Aruba** | 105 | 3 | 2.9 | 2 | 1.9 |
| **AFG** | **Afghanistan** | 445 | 30 | 6.7 | 24 | 5.4 |
| **AGO** | **Angola** | 939 | 161 | 17.1 | 92 | 9.8 |
| **AIA** | **Anguilla** | 106 | 5 | 4.7 | 3 | 2.8 |
| **ALB** | **Albania** | 286 | 7 | 2.4 | 9 | 3.1 |
| **AND** | **Andorra** | 156 | 5 | 3.2 | 4 | 2.6 |
| **ARE** | **United Arab Emirates** | 245 | 19 | 7.8 | 15 | 6.1 |
| **ARG** | **Argentina** | 998 | 94 | 9.4 | 31 | 3.1 |
| **ARM** | **Armenia** | 290 | 10 | 3.4 | 8 | 2.8 |
| **ASM** | **American Samoa** | 69 | 8 | 11.6 | 4 | 5.8 |
| **ATG** | **Antigua and Barbuda** | 130 | 7 | 5.4 | 4 | 3.1 |
| **AUS** | **Australia** | 771 | 87 | 11.3 | 27 | 3.5 |
| **AUT** | **Austria** | 253 | 3 | 1.2 | 4 | 1.6 |
| **BDI** | **Burundi** | 664 | 108 | 16.3 | 57 | 8.6 |
| **BEL** | **Belgium** | 251 | 5 | 2 | 5 | 2 |
| **BEN** | **Benin** | 592 | 58 | 9.8 | 39 | 6.6 |
| **BES** | **Bonaire, Sint Eustatius and Saba** | 142 | 5 | 3.5 | 2 | 1.4 |
| **BFA** | **Burkina Faso** | 492 | 44 | 8.9 | 31 | 6.3 |
| **BGD** | **Bangladesh** | 651 | 52 | 8 | 22 | 3.4 |
| **BGR** | **Bulgaria** | 315 | 7 | 2.2 | 7 | 2.2 |
| **BHS** | **Bahamas** | 249 | 15 | 6 | 11 | 4.4 |
| **BIH** | **Bosnia and Herzegovina** | 278 | 7 | 2.5 | 8 | 2.9 |
| **BLM** | **Saint Barthélemy** | 113 | 5 | 4.4 | 3 | 2.7 |
| **BLR** | **Belarus** | 238 | 1 | 0.4 | 3 | 1.3 |
| **BLZ** | **Belize** | 515 | 42 | 8.2 | 16 | 3.1 |
| **BMU** | **Bermuda** | 46 | 4 | 8.7 | 4 | 8.7 |
| **BOL** | **Bolivia** | 1449 | 172 | 11.9 | 46 | 3.2 |
| **BRA** | **Brazil** | 1835 | 225 | 12.3 | 77 | 4.2 |
| **BRB** | **Barbados** | 148 | 6 | 4.1 | 4 | 2.7 |
| **BRN** | **Brunei Darussalam** | 468 | 50 | 10.7 | 28 | 6 |
| **BTN** | **Bhutan** | 675 | 69 | 10.2 | 27 | 4 |
| **BWA** | **Botswana** | 551 | 56 | 10.2 | 36 | 6.5 |
| **CAF** | **Central African Republic** | 743 | 116 | 15.6 | 62 | 8.3 |
| **CAN** | **Canada** | 477 | 53 | 11.1 | 55 | 11.5 |
| **CCK** | **Cocos (Keeling) Islands** | 24 | 3 | 12.5 | 3 | 12.5 |
| **CHE** | **Switzerland** | 248 | 4 | 1.6 | 5 | 2 |
| **CHL** | **Chile** | 422 | 44 | 10.4 | 16 | 3.8 |
| **CHN** | **China** | 1317 | 182 | 13.8 | 77 | 5.8 |
| **CIV** | **Cote d'Ivoire** | 694 | 97 | 14 | 62 | 8.9 |
| **CMR** | **Cameroon** | 921 | 151 | 16.4 | 87 | 9.4 |
| **COD** | **Congo, The Democratic Republic of** | 1154 | 222 | 19.2 | 133 | 11.5 |
| **COG** | **Congo** | 670 | 112 | 16.7 | 62 | 9.3 |
| **COK** | **Cook Islands** | 64 | 8 | 12.5 | 5 | 7.8 |
| **COL** | **Colombia** | 1925 | 270 | 14 | 60 | 3.1 |
| **COM** | **Comoros** | 96 | 19 | 19.8 | 8 | 8.3 |
| **CPV** | **Cape Verde** | 71 | 6 | 8.5 | 8 | 11.3 |
| **CRI** | **Costa Rica** | 811 | 97 | 12 | 25 | 3.1 |
| **CUB** | **Cuba** | 285 | 28 | 9.8 | 13 | 4.6 |
| **CUW** | **Curaçao** | 114 | 5 | 4.4 | 2 | 1.8 |
| **CXR** | **Christmas Island** | 48 | 6 | 12.5 | 4 | 8.3 |
| **CYM** | **Cayman Islands** | 184 | 8 | 4.3 | 7 | 3.8 |
| **CYP** | **Cyprus** | 208 | 9 | 4.3 | 11 | 5.3 |
| **CZE** | **Czech Republic** | 230 | 4 | 1.7 | 4 | 1.7 |
| **DEU** | **Germany** | 297 | 4 | 1.3 | 4 | 1.3 |
| **DJI** | **Djibouti** | 350 | 23 | 6.6 | 21 | 6 |
| **DMA** | **Dominica** | 138 | 8 | 5.8 | 3 | 2.2 |
| **DNK** | **Denmark** | 240 | 2 | 0.8 | 2 | 0.8 |
| **DOM** | **Dominican Republic** | 232 | 13 | 5.6 | 6 | 2.6 |
| **DZA** | **Algeria** | 316 | 20 | 6.3 | 21 | 6.6 |
| **ECU** | **Ecuador** | 1631 | 221 | 13.5 | 52 | 3.2 |
| **EGY** | **Egypt** | 355 | 24 | 6.8 | 21 | 5.9 |
| **ERI** | **Eritrea** | 568 | 42 | 7.4 | 38 | 6.7 |
| **ESH** | **Western Sahara** | 219 | 14 | 6.4 | 19 | 8.7 |
| **ESP** | **Spain** | 343 | 19 | 5.5 | 21 | 6.1 |
| **EST** | **Estonia** | 241 | 2 | 0.8 | 4 | 1.7 |
| **ETH** | **Ethiopia** | 841 | 96 | 11.4 | 58 | 6.9 |
| **FIN** | **Finland** | 249 | 2 | 0.8 | 4 | 1.6 |
| **FJI** | **Fiji** | 80 | 10 | 12.5 | 4 | 5 |
| **FRA** | **France** | 520 | 52 | 10 | 27 | 5.2 |
| **FRO** | **Faroe Islands** | 65 | 1 | 1.5 | 2 | 3.1 |
| **FSM** | **Micronesia, Federated States of** | 116 | 16 | 13.8 | 6 | 5.2 |
| **GAB** | **Gabon** | 595 | 99 | 16.6 | 52 | 8.7 |
| **GAM** | **Gambia** | 503 | 43 | 8.5 | 35 | 7 |
| **GBR** | **Great_Britain** | 286 | 11 | 3.8 | 13 | 4.5 |
| **GEO** | **Georgia** | 294 | 12 | 4.1 | 11 | 3.7 |
| **GHA** | **Ghana** | 699 | 90 | 12.9 | 59 | 8.4 |
| **GIN** | **Guinea** | 700 | 90 | 12.9 | 61 | 8.7 |
| **GLP** | **Guadeloupe** | 153 | 9 | 5.9 | 3 | 2 |
| **GNB** | **Guinea-Bissau** | 469 | 41 | 8.7 | 33 | 7 |
| **GNQ** | **Equatorial Guinea** | 486 | 86 | 17.7 | 47 | 9.7 |
| **GRC** | **Greece** | 313 | 11 | 3.5 | 12 | 3.8 |
| **GRD** | **Grenada** | 124 | 7 | 5.6 | 4 | 3.2 |
| **GRL** | **Greenland** | 64 | 4 | 6.2 | 3 | 4.7 |
| **GTM** | **Guatemala** | 689 | 72 | 10.4 | 31 | 4.5 |
| **GUF** | **French Guiana** | 686 | 32 | 4.7 | 10 | 1.5 |
| **GUM** | **Guam** | 83 | 11 | 13.3 | 3 | 3.6 |
| **GUY** | **Guyana** | 831 | 56 | 6.7 | 15 | 1.8 |
| **HMD** | **Heard and Mc Donald Islands** | 44 | 7 | 15.9 | 1 | 2.3 |
| **HRV** | **Croatia** | 293 | 8 | 2.7 | 9 | 3.1 |
| **HTI** | **Haiti** | 176 | 8 | 4.5 | 4 | 2.3 |
| **HUN** | **Hungary** | 245 | 3 | 1.2 | 3 | 1.2 |
| **IDN** | **Indonesia** | 1721 | 288 | 16.7 | 84 | 4.9 |
| **IND** | **India** | 1233 | 149 | 12.1 | 64 | 5.2 |
| **IRL** | **Ireland** | 189 | 3 | 1.6 | 5 | 2.6 |
| **IRN** | **Iran, Islamic Republic of** | 458 | 22 | 4.8 | 21 | 4.6 |
| **IRQ** | **Iraq** | 354 | 17 | 4.8 | 13 | 3.7 |
| **ISL** | **Iceland** | 87 | 1 | 1.1 | 3 | 3.4 |
| **ISR** | **Israel** | 342 | 19 | 5.6 | 18 | 5.3 |
| **ITA** | **Italy** | 337 | 10 | 3 | 12 | 3.6 |
| **JAM** | **Jamaica** | 232 | 11 | 4.7 | 7 | 3 |
| **JEY** | **Jersey** | 144 | 3 | 2.1 | 4 | 2.8 |
| **JOR** | **Jordan** | 303 | 19 | 6.3 | 14 | 4.6 |
| **JPN** | **Japan** | 389 | 42 | 10.8 | 27 | 6.9 |
| **KAZ** | **Kazakstan** | 422 | 26 | 6.2 | 24 | 5.7 |
| **KEN** | **Kenya** | 1080 | 180 | 16.7 | 96 | 8.9 |
| **KGZ** | **Kyrgyzstan** | 324 | 19 | 5.9 | 18 | 5.6 |
| **KHM** | **Cambodia** | 559 | 64 | 11.4 | 21 | 3.8 |
| **KIR** | **Kiribati** | 62 | 10 | 16.1 | 9 | 14.5 |
| **KNA** | **Saint Kitts & Nevis** | 110 | 6 | 5.5 | 3 | 2.7 |
| **KOR** | **Korea, Republic of** | 284 | 17 | 6 | 16 | 5.6 |
| **LAO** | **Lao, People's Democratic Republic** | 729 | 94 | 12.9 | 28 | 3.8 |
| **LBR** | **Liberia** | 560 | 77 | 13.8 | 49 | 8.8 |
| **LBY** | **Libyan Arab Jamahiriya** | 235 | 15 | 6.4 | 18 | 7.7 |
| **LCA** | **Saint Lucia** | 127 | 8 | 6.3 | 3 | 2.4 |
| **LIE** | **Liechtenstein** | 161 | 3 | 1.9 | 3 | 1.9 |
| **LKA** | **Sri Lanka** | 356 | 31 | 8.7 | 20 | 5.6 |
| **LSO** | **Lesotho** | 346 | 25 | 7.2 | 14 | 4 |
| **LTU** | **Lithuania** | 238 | 2 | 0.8 | 3 | 1.3 |
| **LUX** | **Luxembourg** | 176 | 2 | 1.1 | 2 | 1.1 |
| **LVA** | **Latvia** | 244 | 2 | 0.8 | 4 | 1.6 |
| **MAF** | **Saint Martin** | 104 | 5 | 4.8 | 3 | 2.9 |
| **MAR** | **Morocco** | 341 | 22 | 6.5 | 25 | 7.3 |
| **MDA** | **Moldova, Republic of** | 253 | 2 | 0.8 | 3 | 1.2 |
| **MDG** | **Madagascar** | 247 | 51 | 20.6 | 24 | 9.7 |
| **MDV** | **Maldives** | 65 | 10 | 15.4 | 7 | 10.8 |
| **MEX** | **Mexico** | 1069 | 156 | 14.6 | 87 | 8.1 |
| **MHL** | **Marshall Islands** | 62 | 7 | 11.3 | 7 | 11.3 |
| **MKD** | **Macedonia, The Former Yugoslav Republic Of** | 266 | 5 | 1.9 | 6 | 2.3 |
| **MLI** | **Mali** | 559 | 46 | 8.2 | 35 | 6.3 |
| **MLT** | **Malta** | 93 | 4 | 4.3 | 4 | 4.3 |
| **MMR** | **Myanmar** | 1067 | 128 | 12 | 58 | 5.4 |
| **MNE** | **Montenegro** | 274 | 6 | 2.2 | 7 | 2.6 |
| **MNG** | **Mongolia** | 370 | 26 | 7 | 21 | 5.7 |
| **MNP** | **Northern Mariana Islands** | 91 | 14 | 15.4 | 4 | 4.4 |
| **MOZ** | **Mozambique** | 731 | 106 | 14.5 | 55 | 7.5 |
| **MRT** | **Mauritania** | 487 | 39 | 8 | 37 | 7.6 |
| **MSR** | **Montserrat** | 103 | 7 | 6.8 | 2 | 1.9 |
| **MTQ** | **Martinique** | 142 | 5 | 3.5 | 2 | 1.4 |
| **MUS** | **Mauritius** | 90 | 15 | 16.7 | 4 | 4.4 |
| **MWI** | **Malawi** | 649 | 95 | 14.6 | 51 | 7.9 |
| **MYS** | **Malaysia** | 595 | 59 | 9.9 | 23 | 3.9 |
| **NAM** | **Namibia** | 615 | 73 | 11.9 | 49 | 8 |
| **NCL** | **New Caledonia** | 155 | 14 | 9 | 4 | 2.6 |
| **NER** | **Niger** | 498 | 41 | 8.2 | 31 | 6.2 |
| **NFK** | **Norfolk Island** | 60 | 8 | 13.3 | 3 | 5 |
| **NGA** | **Nigeria** | 891 | 130 | 14.6 | 76 | 8.5 |
| **NIC** | **Nicaragua** | 694 | 71 | 10.2 | 27 | 3.9 |
| **NLD** | **Netherlands** | 256 | 4 | 1.6 | 4 | 1.6 |
| **NOR** | **Norway** | 258 | 5 | 1.9 | 4 | 1.6 |
| **NPL** | **Nepal** | 804 | 72 | 9 | 32 | 4 |
| **NRU** | **Nauru** | 45 | 6 | 13.3 | 3 | 6.7 |
| **NZL** | **New Zealand** | 266 | 37 | 13.9 | 12 | 4.5 |
| **OMN** | **Oman** | 274 | 26 | 9.5 | 21 | 7.7 |
| **PAK** | **Pakistan** | 650 | 47 | 7.2 | 31 | 4.8 |
| **PAN** | **Panama** | 897 | 106 | 11.8 | 22 | 2.5 |
| **PER** | **Peru** | 1858 | 285 | 15.3 | 65 | 3.5 |
| **PHL** | **Philippines** | 568 | 89 | 15.7 | 15 | 2.6 |
| **PNG** | **Papua New Guinea** | 767 | 146 | 19 | 57 | 7.4 |
| **POL** | **Poland** | 265 | 3 | 1.1 | 3 | 1.1 |
| **PRI** | **Puerto Rico** | 240 | 15 | 6.2 | 7 | 2.9 |
| **PRK** | **Korea, Democratic People's Republic of** | 309 | 23 | 7.4 | 20 | 6.5 |
| **PRT** | **Portugal** | 301 | 19 | 6.3 | 19 | 6.3 |
| **PRY** | **Paraguay** | 694 | 43 | 6.2 | 13 | 1.9 |
| **PYF** | **French Polynesia** | 121 | 18 | 14.9 | 9 | 7.4 |
| **QAT** | **Qatar** | 158 | 12 | 7.6 | 9 | 5.7 |
| **ROU** | **Romania** | 295 | 4 | 1.4 | 5 | 1.7 |
| **RUS** | **Russia** | 675 | 52 | 7.7 | 45 | 6.7 |
| **RWA** | **Rwanda** | 708 | 117 | 16.5 | 64 | 9 |
| **SAU** | **Saudi Arabia** | 354 | 33 | 9.3 | 22 | 6.2 |
| **SEN** | **Senegal** | 584 | 49 | 8.4 | 41 | 7 |
| **SGP** | **Singapore** | 366 | 32 | 8.7 | 13 | 3.6 |
| **SGS** | **South Georgia & The South Sandwich Islands** | 46 | 4 | 8.7 | 1 | 2.2 |
| **SHN** | **Saint Helena** | 109 | 10 | 9.2 | 7 | 6.4 |
| **SLB** | **Solomon Islands** | 246 | 44 | 17.9 | 10 | 4.1 |
| **SLE** | **Sierra Leone** | 608 | 81 | 13.3 | 51 | 8.4 |
| **SOM** | **Somalia** | 619 | 87 | 14.1 | 51 | 8.2 |
| **SPM** | **Saint Pierre and Miquelon** | 113 | 3 | 2.7 | 5 | 4.4 |
| **SRB** | **Republic of Serbia** | 275 | 4 | 1.5 | 6 | 2.2 |
| **STP** | **Sao Tome and Principe** | 97 | 13 | 13.4 | 5 | 5.2 |
| **SUR** | **Suriname** | 706 | 34 | 4.8 | 13 | 1.8 |
| **SVK** | **Slovakia** | 238 | 3 | 1.3 | 3 | 1.3 |
| **SVN** | **Slovenia** | 266 | 5 | 1.9 | 5 | 1.9 |
| **SWE** | **Sweden** | 265 | 4 | 1.5 | 4 | 1.5 |
| **SWZ** | **Swaziland** | 508 | 49 | 9.6 | 23 | 4.5 |
| **SYC** | **Seychelles** | 90 | 16 | 17.8 | 6 | 6.7 |
| **TCD** | **Chad** | 561 | 53 | 9.4 | 36 | 6.4 |
| **TGO** | **Togo** | 608 | 62 | 10.2 | 43 | 7.1 |
| **THA** | **Thailand** | 922 | 108 | 11.7 | 43 | 4.7 |
| **TJK** | **Tajikistan** | 323 | 19 | 5.9 | 15 | 4.6 |
| **TMP** | **Timor Leste** | 241 | 18 | 7.5 | 8 | 3.3 |
| **TON** | **Tonga** | 77 | 11 | 14.3 | 4 | 5.2 |
| **TUN** | **Tunisia** | 295 | 16 | 5.4 | 18 | 6.1 |
| **TUR** | **Turkey** | 372 | 18 | 4.8 | 17 | 4.6 |
| **TUV** | **Tuvalu** | 51 | 6 | 11.8 | 5 | 9.8 |
| **TWN** | **Taiwan, Province of China** | 335 | 33 | 9.9 | 16 | 4.8 |
| **TZA** | **Tanzania** | 1101 | 203 | 18.4 | 103 | 9.4 |
| **UGA** | **Uganda** | 1026 | 174 | 17 | 91 | 8.9 |
| **UKR** | **Ukraine** | 306 | 5 | 1.6 | 6 | 2 |
| **URY** | **Uruguay** | 424 | 27 | 6.4 | 13 | 3.1 |
| **USA** | **USA** | 906 | 118 | 13 | 95 | 10.5 |
| **VCT** | **Saint Vincent and the Grenadines** | 123 | 7 | 5.7 | 3 | 2.4 |
| **VEN** | **Venezuela** | 1402 | 143 | 10.2 | 28 | 2 |
| **VGB** | **Virgin Islands, British** | 142 | 6 | 4.2 | 4 | 2.8 |
| **VNM** | **Vietnam** | 829 | 114 | 13.8 | 35 | 4.2 |
| **VUT** | **Vanuatu** | 124 | 13 | 10.5 | 5 | 4 |
| **WLF** | **Wallis and Futuna** | 58 | 6 | 10.3 | 4 | 6.9 |
| **WSM** | **Samoa** | 85 | 9 | 10.6 | 4 | 4.7 |
| **YEM** | **Yemen** | 282 | 30 | 10.6 | 21 | 7.4 |
| **ZAF** | **South Africa** | 762 | 90 | 11.8 | 47 | 6.2 |
| **ZMB** | **Zambia** | 744 | 114 | 15.3 | 61 | 8.2 |
| **ZWE** | **Zimbabwe** | 632 | 76 | 12 | 39 | 6.2 |

**Table S2: Number of eponyms and proportion of all eponyms attributed to persons belonging to each country.**

| **Nationalities of persons named in eponyms worldwide** | **Number of scientific name eponyms attributed to the country** | **Percentage of all scientific name eponyms that honor a person from the country** | **Number of common name eponyms attributed to the country** | **Percentage of all common name eponyms that honor a person from the country** |
| --- | --- | --- | --- | --- |
| **Argentina** | 9 | 0.41 | 4 | 0.54 |
| **Australia** | 49 | 2.22 | 9 | 1.21 |
| **Austria** | 39 | 1.76 | 11 | 1.48 |
| **Belgium** | 16 | 0.72 | 8 | 1.08 |
| **Bolivia** | 2 | 0.09 | 1 | 0.13 |
| **Brazil** | 24 | 1.09 | 6 | 0.81 |
| **Bulgaria** | 1 | 0.05 | 0 | 0 |
| **Cambodia** | 1 | 0.05 | 0 | 0 |
| **Canada** | 4 | 0.18 | 3 | 0.4 |
| **Chile** | 1 | 0.05 | 0 | 0 |
| **China** | 1 | 0.05 | 0 | 0 |
| **Colombia** | 14 | 0.63 | 4 | 0.54 |
| **Costa Rica** | 4 | 0.18 | 1 | 0.13 |
| **Cuba** | 8 | 0.36 | 1 | 0.13 |
| **Cyprus** | 1 | 0.05 | 0 | 0 |
| **Czech Republic** | 2 | 0.09 | 2 | 0.27 |
| **Denmark** | 7 | 0.32 | 1 | 0.13 |
| **Ecuador** | 3 | 0.14 | 0 | 0 |
| **Egypt** | 1 | 0.05 | 0 | 0 |
| **Estonia** | 1 | 0.05 | 0 | 0 |
| **Finland** | 1 | 0.05 | 0 | 0 |
| **France** | 348 | 15.75 | 59 | 7.94 |
| **Germany** | 323 | 14.62 | 116 | 15.61 |
| **Great_Britain** | 647 | 29.28 | 244 | 32.84 |
| **Greece** | 2 | 0.09 | 0 | 0 |
| **Guadeloupe** | 2 | 0.09 | 0 | 0 |
| **Guatemala** | 1 | 0.05 | 0 | 0 |
| **Hungary** | 6 | 0.27 | 2 | 0.27 |
| **India** | 2 | 0.09 | 0 | 0 |
| **Indonesia** | 3 | 0.14 | 0 | 0 |
| **Ireland** | 10 | 0.45 | 3 | 0.4 |
| **Israel** | 1 | 0.05 | 0 | 0 |
| **Italy** | 42 | 1.9 | 23 | 3.1 |
| **Japan** | 11 | 0.5 | 3 | 0.4 |
| **Latvia** | 1 | 0.05 | 0 | 0 |
| **Liberia** | 1 | 0.05 | 0 | 0 |
| **Luxembourg** | 1 | 0.05 | 0 | 0 |
| **Madagascar** | 1 | 0.05 | 0 | 0 |
| **Mauritius** | 1 | 0.05 | 0 | 0 |
| **Mexico** | 9 | 0.41 | 2 | 0.27 |
| **Mozambique** | 1 | 0.05 | 0 | 0 |
| **Netherlands** | 106 | 4.8 | 17 | 2.29 |
| **New Zealand** | 17 | 0.77 | 4 | 0.54 |
| **Nigeria** | 1 | 0.05 | 0 | 0 |
| **Norway** | 4 | 0.18 | 1 | 0.13 |
| **Peru** | 6 | 0.27 | 3 | 0.4 |
| **Philippines** | 3 | 0.14 | 1 | 0.13 |
| **Poland** | 28 | 1.27 | 7 | 0.94 |
| **Portugal** | 19 | 0.86 | 8 | 1.08 |
| **Russia** | 26 | 1.18 | 22 | 2.96 |
| **Saint Lucia** | 1 | 0.05 | 1 | 0.13 |
| **Sao Tome and Princepe** | 1 | 0.05 | 0 | 0 |
| **South Africa** | 17 | 0.77 | 11 | 1.48 |
| **Spain** | 9 | 0.41 | 3 | 0.4 |
| **Sri Lanka** | 1 | 0.05 | 0 | 0 |
| **Sweden** | 13 | 0.59 | 6 | 0.81 |
| **Switzerland** | 18 | 0.81 | 8 | 1.08 |
| **Tanzania** | 1 | 0.05 | 0 | 0 |
| **Thailand** | 1 | 0.05 | 0 | 0 |
| **Trinidad** | 1 | 0.05 | 1 | 0.13 |
| **Turkey** | 1 | 0.05 | 1 | 0.13 |
| **USA** | 320 | 14.48 | 136 | 18.3 |
| **Venezuela** | 1 | 0.05 | 0 | 0 |

**Table S3: Number of country-matching eponyms found in each modern-day country (where the species is named after a person from the country where it occurs) and the percentage of all eponyms which are country-matching in each country, shown separately for scientific and common names.**

| **Country** | **Number of country-matching scientific name eponyms in the country (in brackets: total number of eponymous species found in the country)** | **Percentage of all scientific name eponyms which are country-matching** | **Number of country-matching common name eponyms in the country (in brackets: total number of eponymous species found in the country)** | **Percentage of all common name eponyms which are country-matching** |
| --- | --- | --- | --- | --- |
| **USA** | 58 (118) | 49.2 | 63 (95) | 66.3 |
| **Australia** | 26 (87) | 29.9 | 5 (27) | 18.5 |
| **Brazil** | 22 (225) | 9.8 | 5 (77) | 6.5 |
| **France** | 15 (52) | 28.8 | 3 (27) | 11.1 |
| **Colombia** | 14 (270) | 5.2 | 4 (60) | 6.7 |
| **New Zealand** | 11 (37) | 29.7 | 4 (12) | 33.3 |
| **South Africa** | 10 (90) | 11.1 | 7 (47) | 14.9 |
| **Argentina** | 8 (94) | 8.5 | 4 (31) | 12.9 |
| **Cuba** | 8 (28) | 28.6 | 1 (13) | 7.7 |
| **Mexico** | 8 (156) | 5.1 | 2 (87) | 2.3 |
| **Japan** | 6 (42) | 14.3 | 3 (27) | 11.1 |
| **Peru** | 6 (285) | 2.1 | 3 (65) | 4.6 |
| **Russia** | 6 (52) | 11.5 | 14 (45) | 31.1 |
| **Costa Rica** | 4 (97) | 4.1 | 1 (25) | 4 |
| **Italy** | 4 (10) | 40 | 8 (12) | 66.7 |
| **Ecuador** | 3 (221) | 1.4 | 0 (52) | 0 |
| **Great Britain** | 3 (11) | 27.3 | 5 (13) | 38.5 |
| **Indonesia** | 3 (288) | 1 | 0 (84) | 0 |
| **Philippines** | 3 (89) | 3.4 | 1 (15) | 6.7 |
| **Bolivia** | 2 (172) | 1.2 | 1 (46) | 2.2 |
| **Guadeloupe** | 2 (9) | 22.2 | 0 (3) | 0 |
| **India** | 2 (149) | 1.3 | 0 (64) | 0 |
| **Cambodia** | 1 (64) | 1.6 | 0 (21) | 0 |
| **Canada** | 1 (53) | 1.9 | 1 (55) | 1.8 |
| **China** | 1 (182) | 0.5 | 0 (77) | 0 |
| **Germany** | 1 (4) | 25 | 0 (4) | 0 |
| **Greece** | 1 (11) | 9.1 | 0 (12) | 0 |
| **Israel** | 1 (19) | 5.3 | 0 (18) | 0 |
| **Liberia** | 1 (77) | 1.3 | 0 (49) | 0 |
| **Madagascar** | 1 (51) | 2 | 0 (24) | 0 |
| **Mauritius** | 1 (15) | 6.7 | 0 (4) | 0 |
| **Nigeria** | 1 (130) | 0.8 | 0 (76) | 0 |
| **Saint Lucia** | 1 (8) | 12.5 | 1 (3) | 33.3 |
| **Sri Lanka** | 1 (31) | 3.2 | 0 (20) | 0 |
| **Tanzania** | 1 (203) | 0.5 | 0 (103) | 0 |
| **Thailand** | 1 (108) | 0.9 | 0 (43) | 0 |
| **Venezuela** | 1 (143) | 0.7 | 0 (28) | 0 |
| **Portugal** | 0 (19) | 0 | 1 (19) | 5.3 |

**Table S4: The reasons/professions of the honorees in all eponyms found worldwide.**

| **Occupation/Reason for naming** | **Total number of eponyms honoring person with the occupation/reason** |
| --- | --- |
| **Scientist** | 892 |
| **Naturalist** | 394 |
| **Civilian** | 253 |
| **Army/Civil Servant** | 200 |
| **Personal relation** | 108 |
| **Royalty/Landowner** | 107 |
| **Businessman** | 94 |
| **Doctor** | 78 |
| **Religious person** | 47 |
| **Artist** | 26 |
| **Conservationist** | 6 |

**Table S5: The order of priority when marking professions of persons named in eponyms, as stated in the Book of Bird Eponyms** **(Beolens et al. 2020)****. This order was needed to distinguish naturalists and collectors from polymaths and professionals defined by the field of study they undertook. For example, to distinguish an amateur collector of bird specimens from an army doctor or a professional zoologist.**

| **Priority** | **Criteria** | **Example** |
| --- | --- | --- |
| 1 | If there is a mention of any professional training, it is placed at highest priority. | Doctor, Lawyer |
| 2 | If there was a field of science that the person was specialized in, it was given the next importance. | Zoologist, botanist |
| 3 | If no other specific profession or field of scientific study was presented, persons were considered only a naturalist or collector. | Naturalist, Collector |

**Table S6: Model estimates of the association between national counts of common name eponyms and colonizer CEI scores. The model includes observation-level random effects and random intercepts for countries and colonizing powers. R-hat and effective sample sizes (ESS) are reported for all parameters. R-hat was 1 with effective sample sizes (ESS) over 5000 for all parameters.**

| **Term** | **Poisson component** | | | **Zero-inflated Component** | | |
| --- | --- | --- | --- | --- | --- | --- |
|  | **Estimate** | **Q2.5** | **Q97.5** | **Estimate** | **Q2.5** | **Q97.5** |
| **Intercept** | -6.90 | -8.51 | -5.4 | -3.86 | -8.49 | 1.74 |
| **Effect of CEI** | 1.2 | 0.37 | 2.16 | -9.87 | -35.92 | 1.06 |
| **colonizer-level random intercepts** | 2.17 | 1.26 | 3.64 | 4.07 | 0.20 | 15.38 |
| **Country-level random intercepts** | 0.60 | 0.50 | 0.71 | 1.06 | 0.03 | 3.38 |
| **Observation-level random effect** | 0.37 | 0.3 | 0.44 |  |  |  |
| **colonizer-level random slopes** | 1.06 | 0.03 | 3.38 |  |  |  |
| **Correlation between colonizer-specific intercepts and slopes** | -0.79 | -0.99 | -0.20 |  |  |  |

**Supplementary Analysis 1: Assigning nationality by birth (not following the decision tree)**

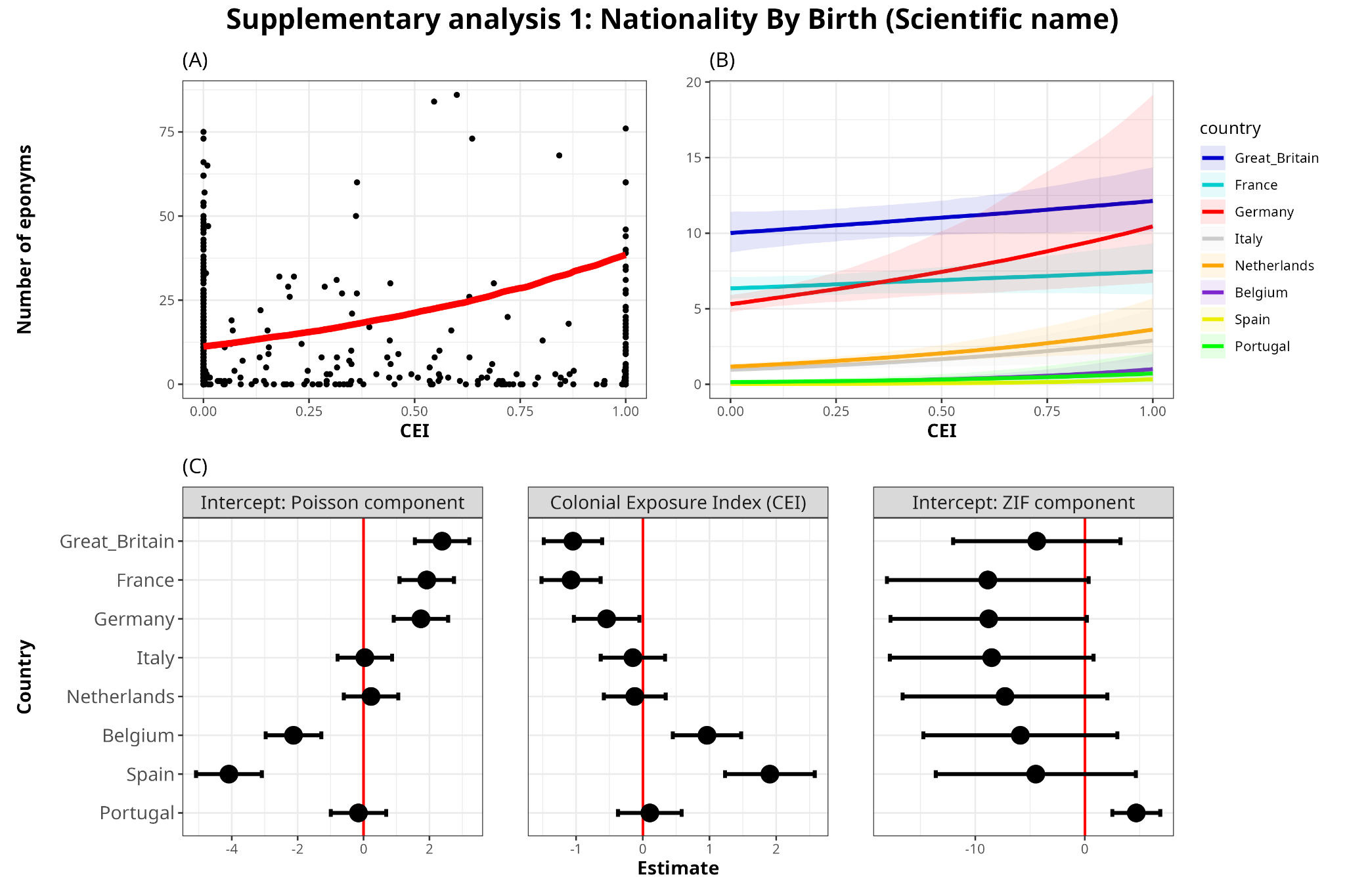

**Figure S6: (A)** Model-predicted relationship between CEI and number of eponyms. Black points represent raw data points, while the red line represents the central tendency of the relationship (posterior median, marginalized over colonizers), when nationality is assigned by country of birth; **(B)** Actual effects of CEI on the number of eponyms (scientific names) for each European colonizer. In the top row, the scale is absolute and represents the actual number of predicted eponyms on the vertical axis. **(C)** Latent scale parameter estimates with the 95% confidence intervals of the Poisson-component intercepts, effect size of Colonial Exposure Index (CEI) and the Zero-inflated-component intercept on the number of eponyms for each colonizer (in scientific names).

**Table S7: Model estimates of the relationship between national counts of scientific name eponyms and colonizers CEI scores for the country – when using nationality assigned by country of birth, not following our decision tree.** The model includes observation-level random effects and random intercepts for countries and colonizing powers. R-hat was 1 with effective sample sizes (ESS) over 5000 for all parameters, indicating good convergence and reliable posterior estimation in the model.

| **Term** | **Effect type** | **Poisson component** | | | **Zero-inflated Component** | | |
| --- | --- | --- | --- | --- | --- | --- | --- |
|  |  | **Estimate** | **Q2.5** | **Q97.5** | **Estimate** | **Q2.5** | **Q97.5** |
| **Intercept** | **Fixed** | -5.97 | -7.63 | -4.3 | -2.99 | -7.36 | 0.99 |
| **CEI** | **Fixed** | 1.24 | 0.42 | 2.13 | -0.39 | -2.69 | 1.79 |
| **colonizer-level intercepts** | **Random** | 2.35 | 1.41 | 3.97 | 8.09 | 2.53 | 25.22 |
| **Country-level intercepts** | **Random** | 0.31 | 0.24 | 0.39 | 1.37 | 0.05 | 4.69 |
| **Observation-level effect** | **Random** | 0.40 | 0.35 | 0.45 |  |  |  |
| **colonizer-level slopes** | **Random** | 1.12 | 0.58 | 2.04 |  |  |  |
| **Correlation between colonizer-specific intercepts and slopes** | **Random** | -0.91 | -1 | -0.57 |  |  |  |

**Supplementary Analysis 2: Excluding migratory species for analyses**

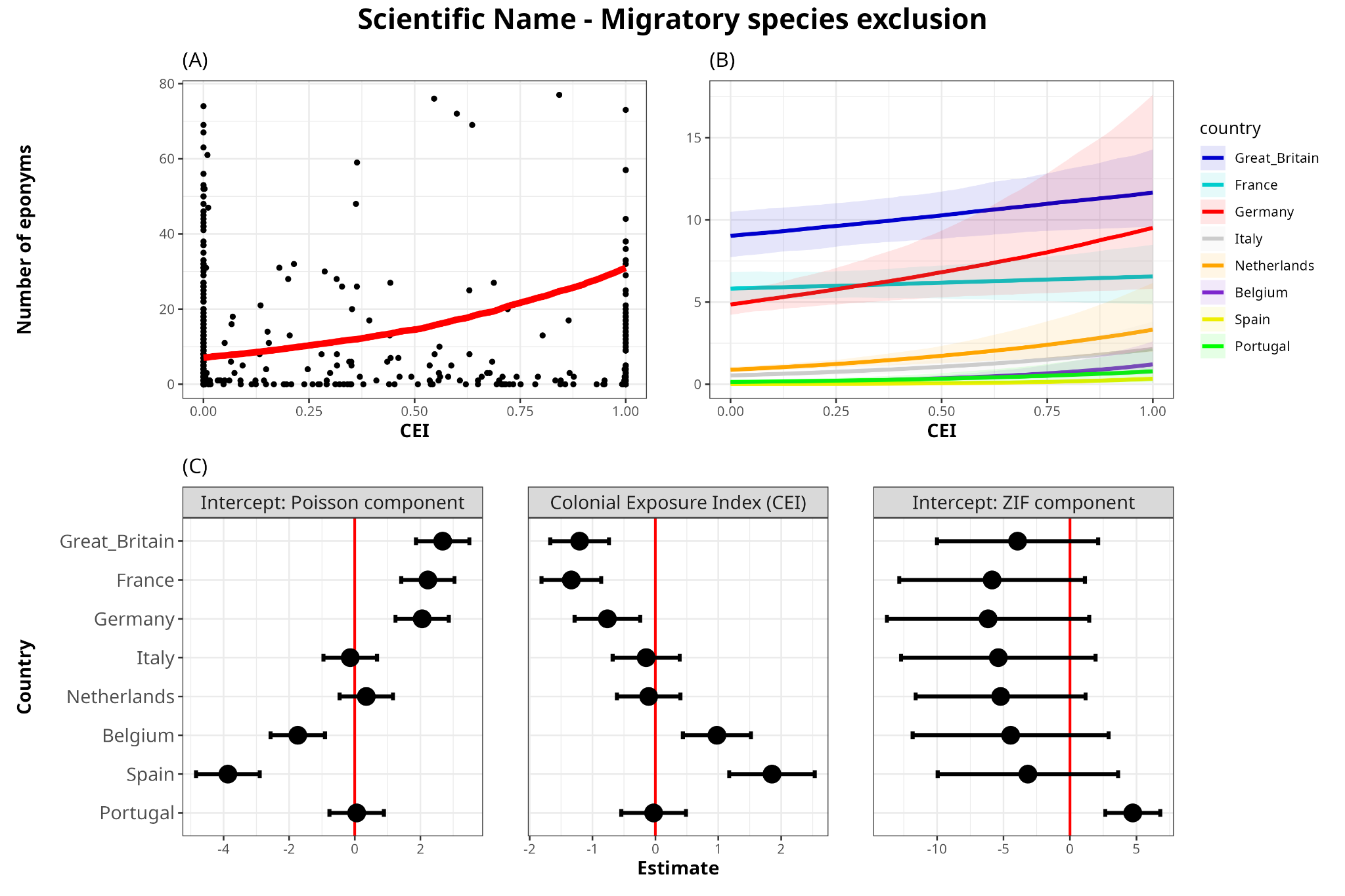

**Figure S7: (A)** Model-predicted relationship between CEI and number of eponyms. Black points represent raw data points, while the red line represents the central tendency of the relationship (posterior median, marginalized over colonizers), when excluding all migratory species; **(B)** Actual effects of CEI on the number of eponyms (scientific names) for each European colonizer. In the top row, the scale is absolute and represents the actual number of predicted eponyms on the vertical axis. **(C)** Latent scale parameter estimates with the 95% confidence intervals of the Poisson-component intercepts, effect size of Colonial Exposure Index (CEI) and the Zero-inflated-component intercept on the number of eponyms for each colonizer (in scientific names).

**Table S8: Model estimates of the relationship between national counts of scientific name eponyms and colonizers CEI scores for the country – when excluding all migratory species from analyses.** The model includes observation-level random effects and random intercepts for modern-day countries and colonial empires. R-hat was 1 with effective sample sizes (ESS) over 5000 for all parameters, indicating good convergence and reliable posterior estimation in the model.

| **Term** | **Effect type** | **Poisson component** | | | **Zero-inflated Component** | | |
| --- | --- | --- | --- | --- | --- | --- | --- |
|  |  | **Estimate** | **Q2.5** | **Q97.5** | **Estimate** | **Q2.5** | **Q97.5** |
| **Intercept** | **Fixed** | -6.48 | -8.16 | -4.94 | -3.33 | -7.26 | 0.48 |
| **CEI** | **Fixed** | 1.46 | 0.63 | 2.42 | -0.41 | -2.60 | 1.68 |
| **Colonizer-level intercepts** | **Random** | 2.30 | 1.41 | 3.82 | 6.06 | 2.07 | 18.19 |
| **Country-level intercepts** | **Random** | 0.53 | 0.44 | 0.64 | 1.20 | 0.05 | 4.00 |
| **Observation-level effect** | **Random** | 0.43 | 0.38 | 0.49 |  |  |  |
| **Colonizer-level slopes** | **Random** | 1.17 | 0.63 | 2.07 |  |  |  |
| **Correlation between colonizer-specific intercepts and slopes** | **Random** | -0.90 | -1 | -0.54 |  |  |  |

**Diagnostic plots for the scientific name model**
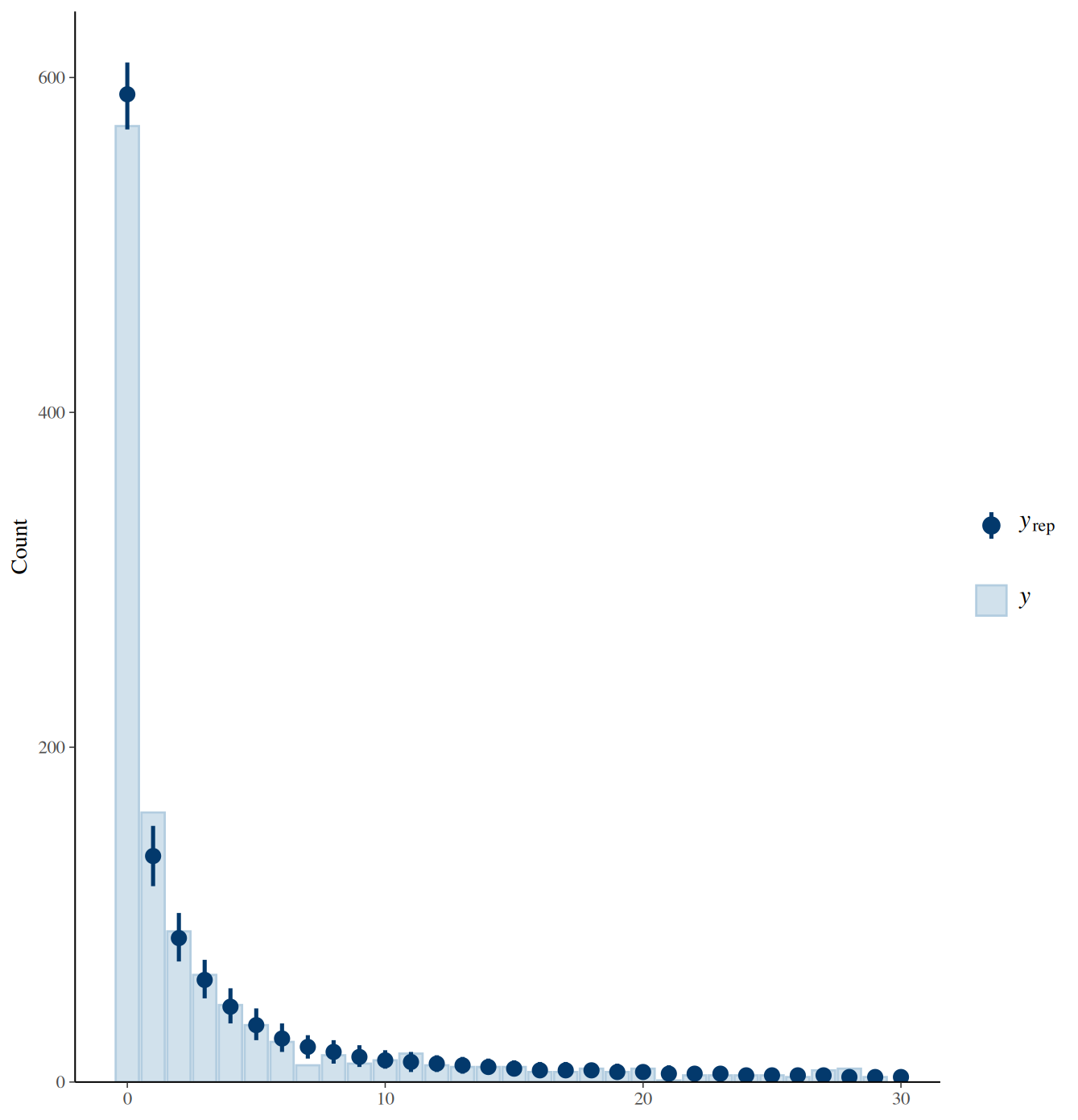
 **(Figure 4):**

**Diagnostic plot #1: Posterior predictive check for the scientific name model. Simulated data (blue) closely match the observed distribution (black), indicating good model fit.**

**Diagnostic plot #2: Trace and density plots for key parameters of the model reported in Figure 6. The plots indicate good mixing and convergence across chains.**

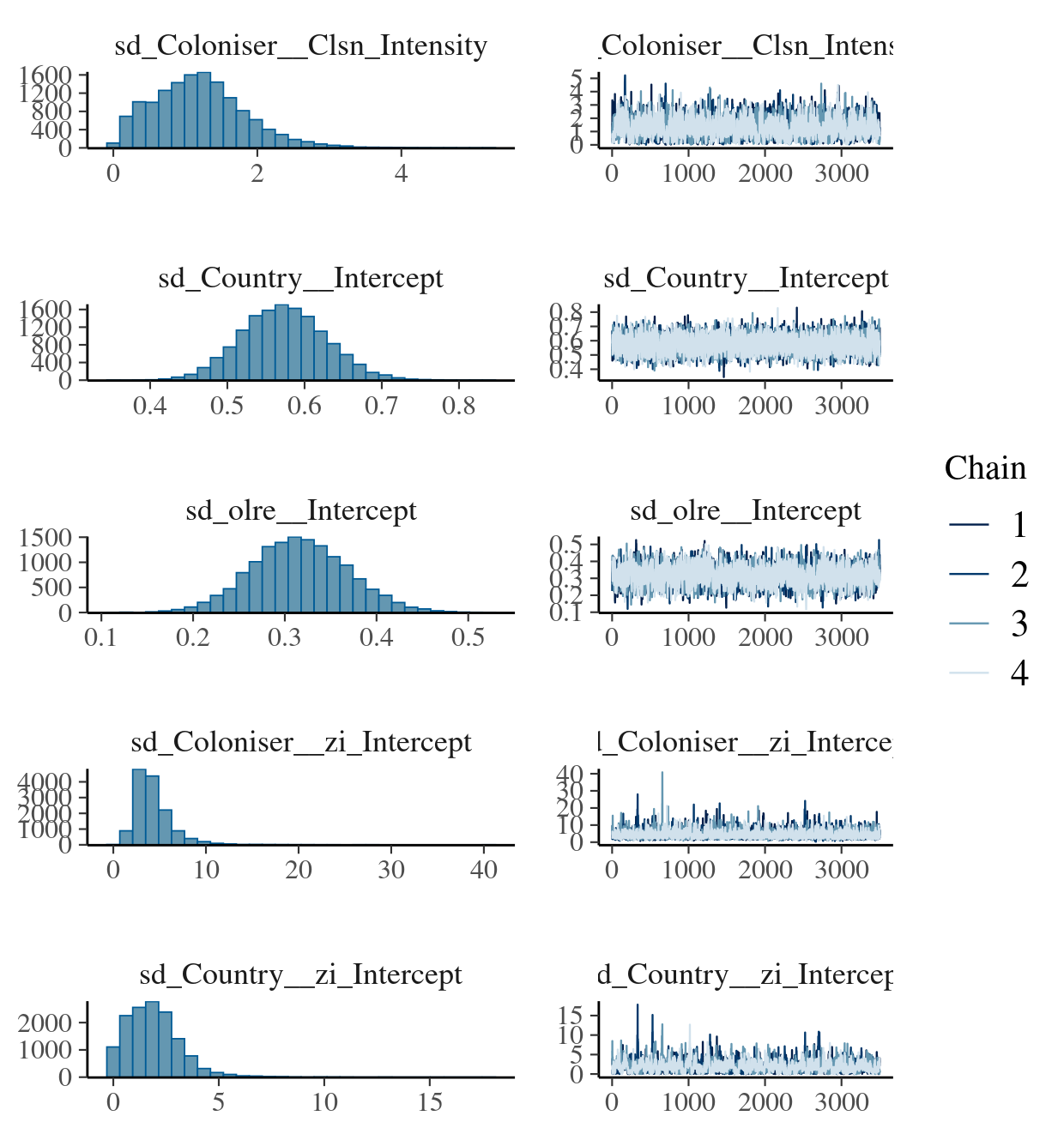

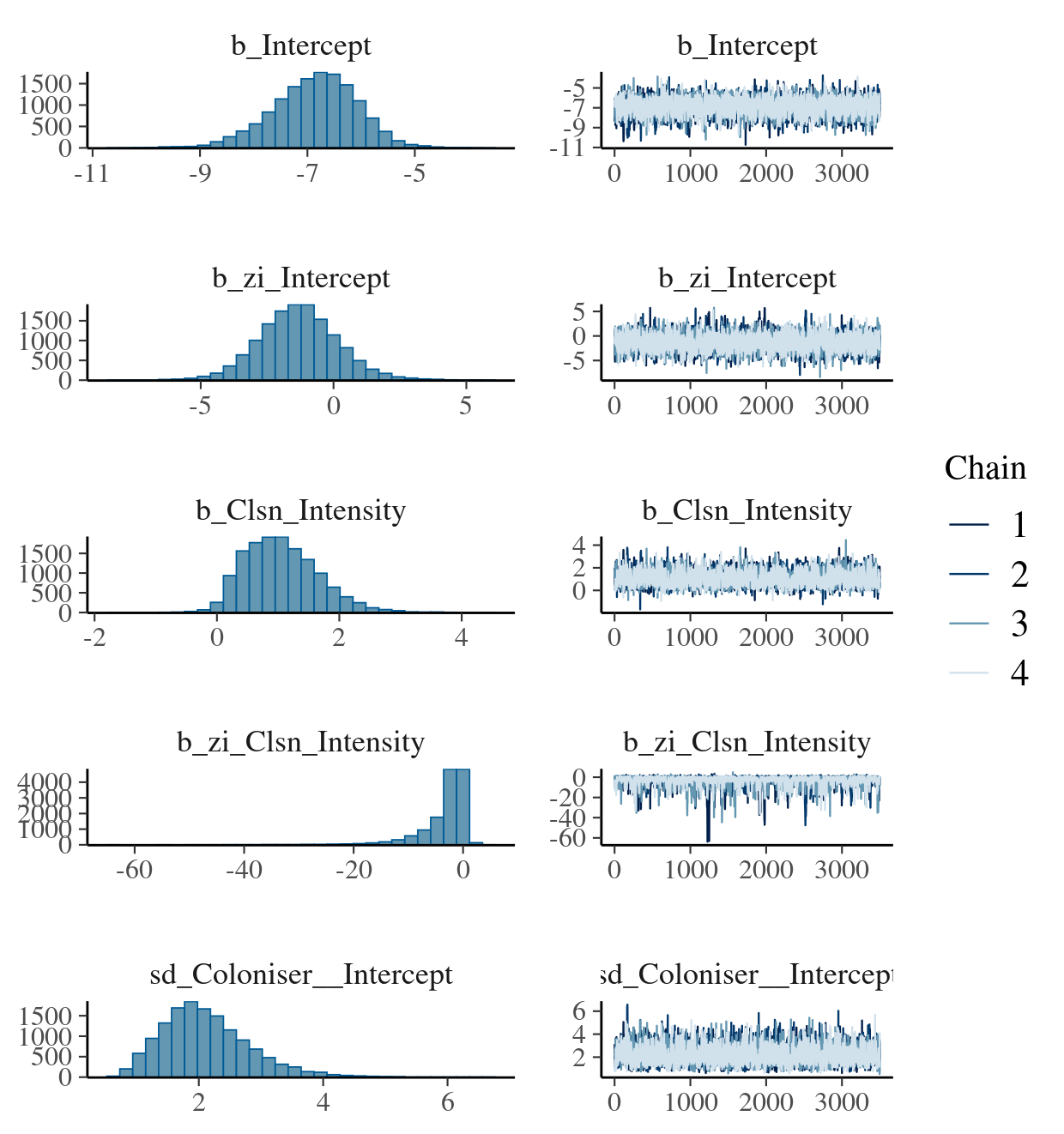

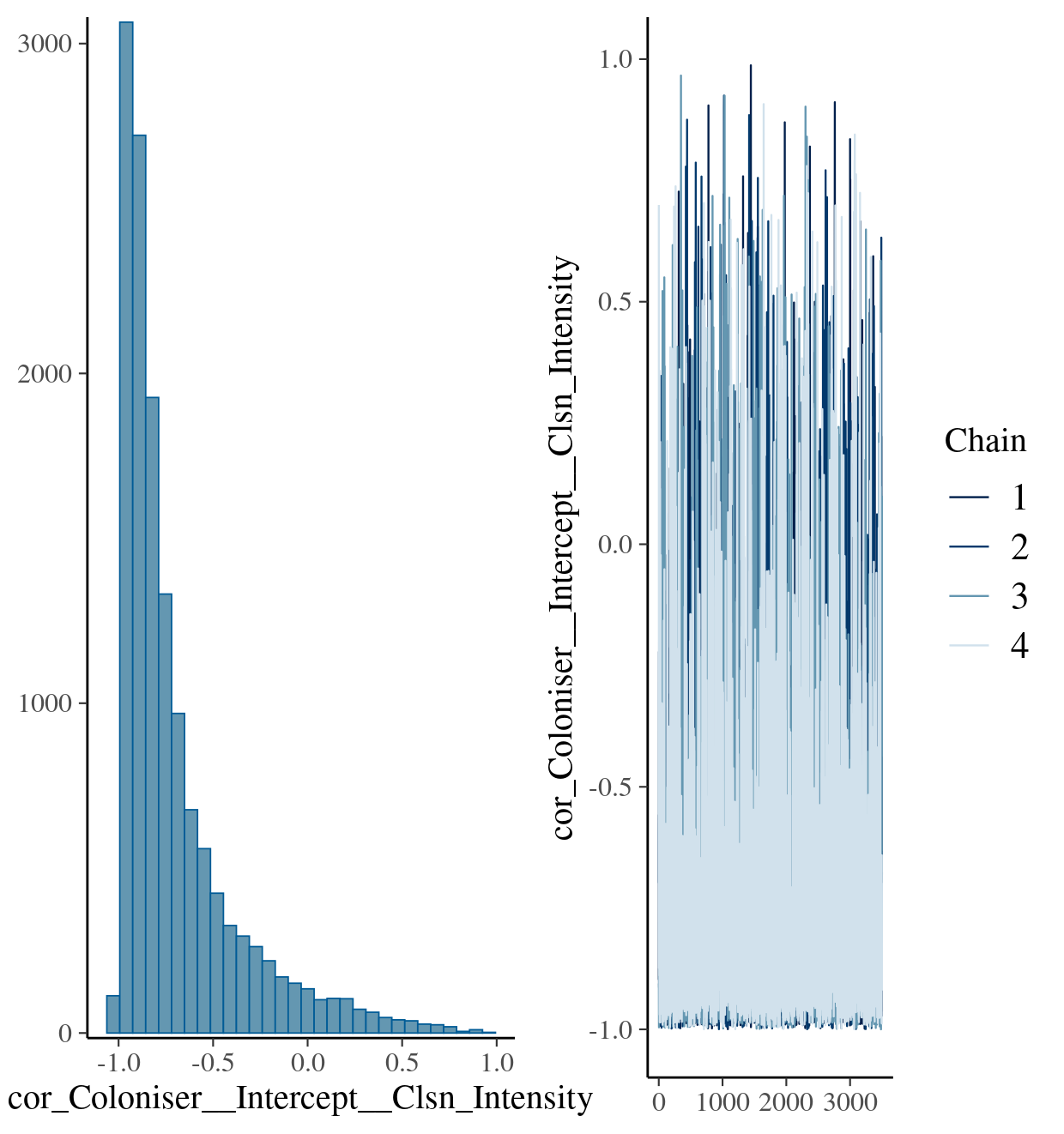

**Diagnostic plots for the common name model**
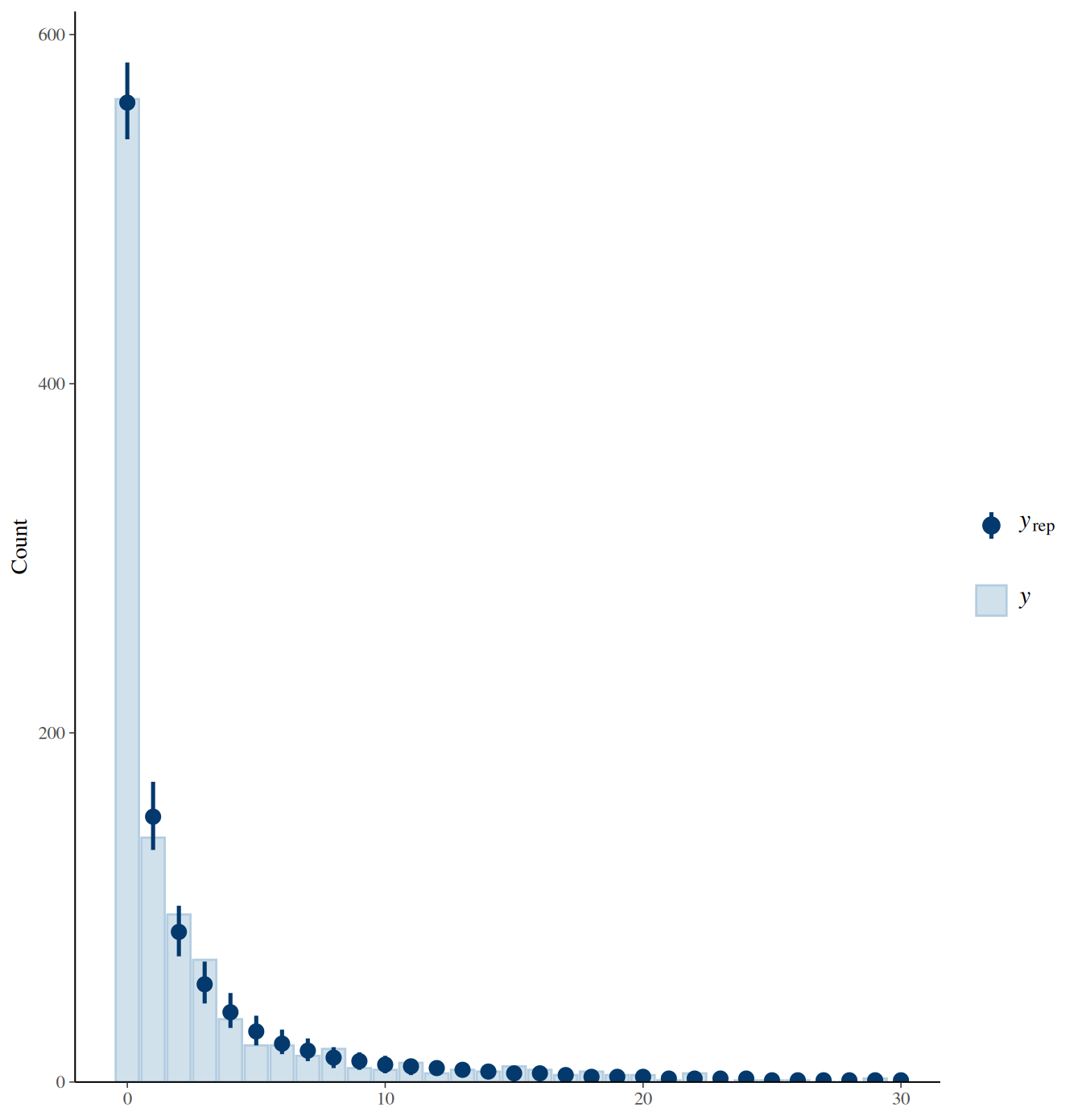
 **(Fig S4):**

**Diagnostic plot #1: Posterior predictive check for the common name model reported in Figure S4. Simulated data (blue) closely match the observed distribution (black), indicating good model fit.**

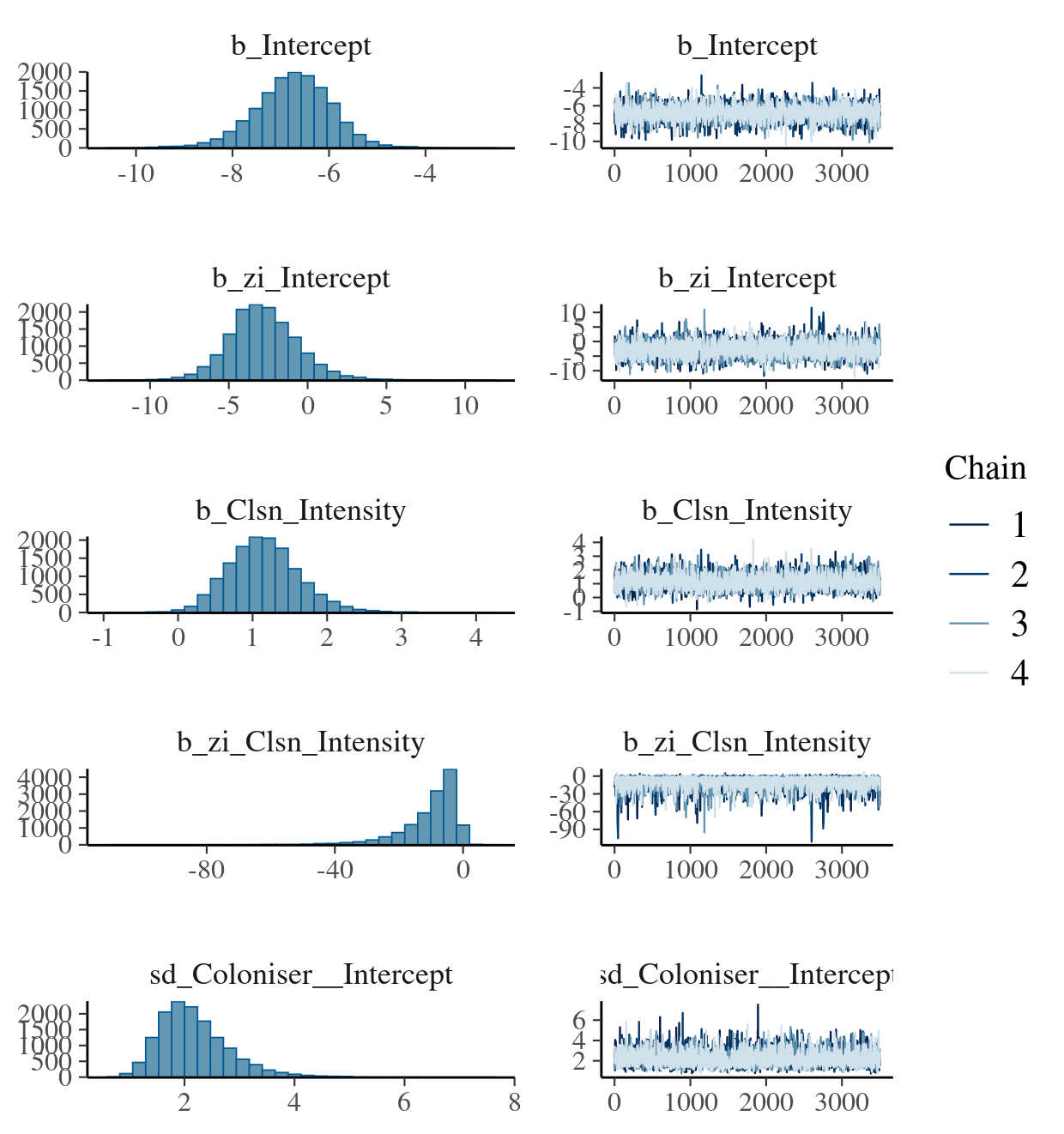

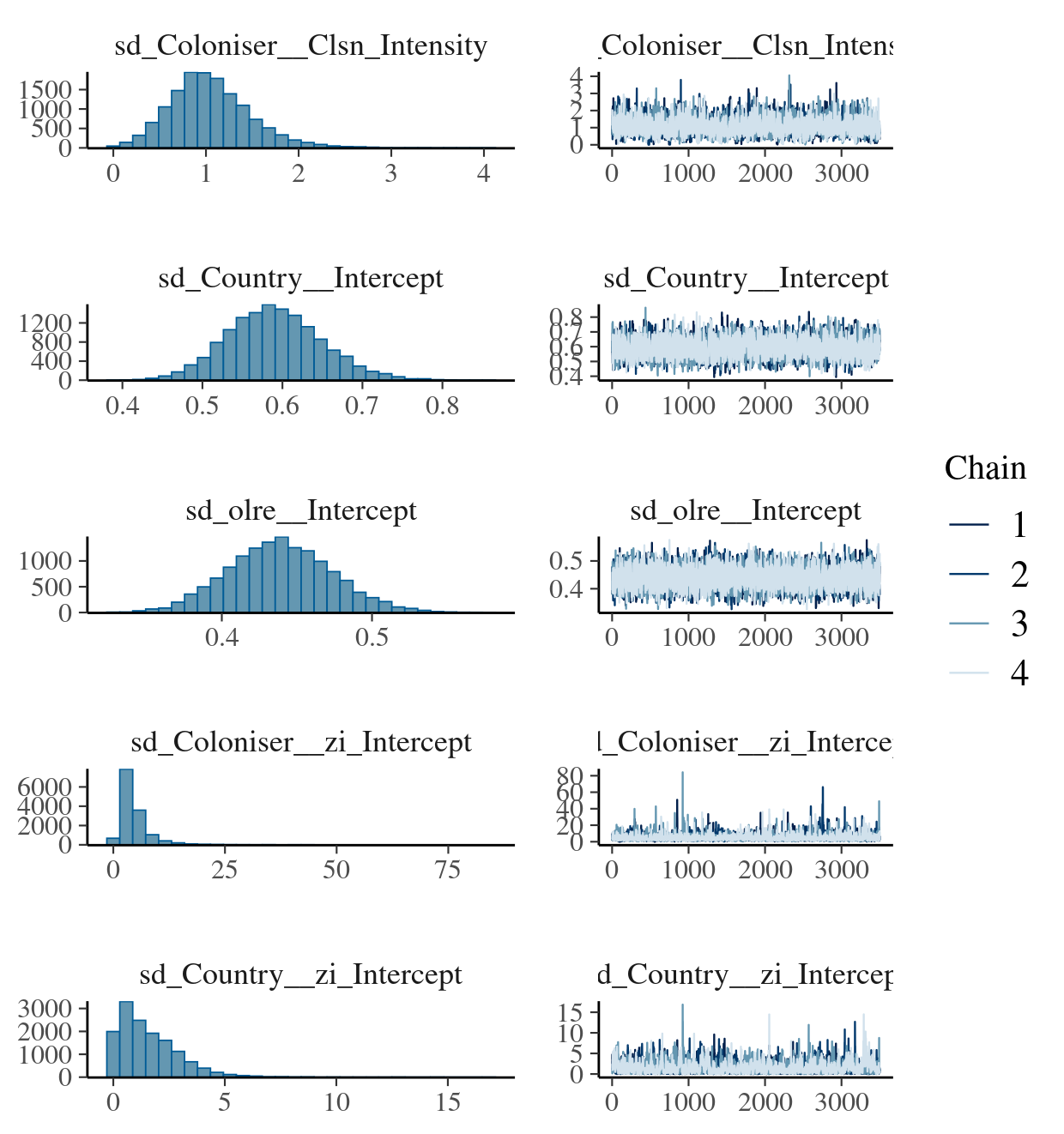

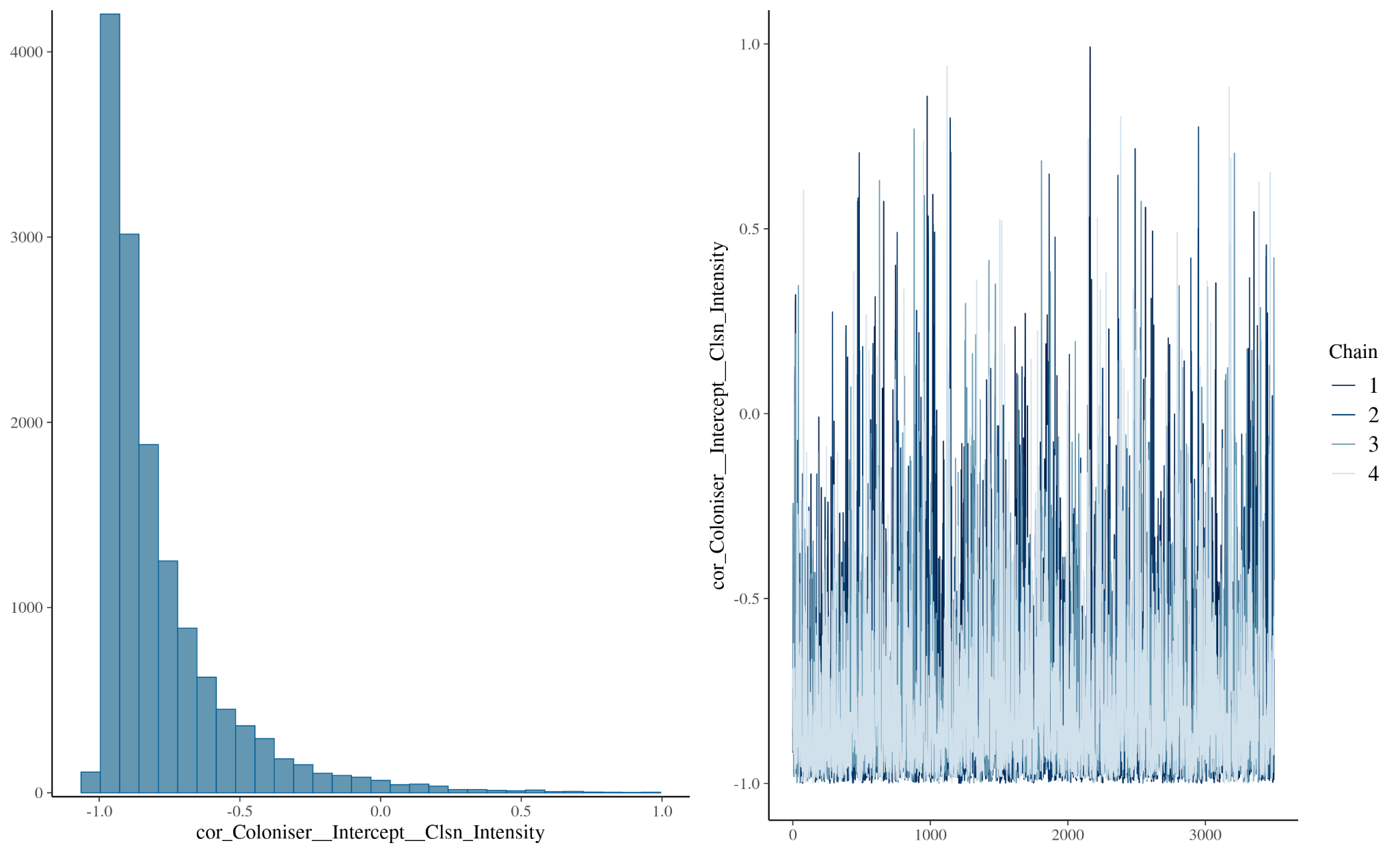
**Diagnostic plot #2: Trace and density plots for key parameters of the model reported in Figure S4. The plots indicate good mixing and convergence across chains.**
